# *Trans* control of cardiac mRNA translation in a protein length-dependent fashion

**DOI:** 10.1101/2020.06.05.133298

**Authors:** Franziska Witte, Jorge Ruiz-Orera, Camilla Ciolli Mattioli, Susanne Blachut, Eleonora Adami, Jana Felicitas Schulz, Valentin Schneider-Lunitz, Oliver Hummel, Giannino Patone, Michael Benedikt Mücke, Jan Šilhavý, Matthias Heinig, Leonardo Bottolo, Daniel Sanchis, Martin Vingron, Marina Chekulaeva, Michal Pravenec, Norbert Hubner, Sebastiaan van Heesch

**Author notes:** These authors contributed equally (Franziska Witte, Jorge Ruiz-Orera). Present address: The Princess Máxima Center for Pediatric Oncology, Utrecht, the Netherlands (Sebastiaan van Heesch).

## Abstract

Little is known about the impact of naturally occurring genetic variation on the rates with which proteins are synthesized by ribosomes. Here, we investigate how genetic influences on mRNA translational efficiency are associated with complex disease phenotypes using a panel of rat recombinant inbred lines. We identify a locus for cardiac hypertrophy that is associated with a translatome-wide and protein length-dependent shift in translational efficiency. This master regulator primarily affects the translation of very short and very long protein-coding sequences, altering the physiological stoichiometric translation rates of sarcomere proteins. Mechanistic dissection of this locus points to altered ribosome assembly, characterized by accumulation of polysome half-mers, changed ribosomal configurations and misregulation of the small nucleolar RNA *SNORA48*. We postulate that this locus enhances a pre-existing negative correlation between protein length and translation initiation in diseased hearts. Our work shows that a single genomic locus can trigger a complex, translation-driven molecular mechanism that contributes to phenotypic variability between individuals.

**Graphical Abstract:** 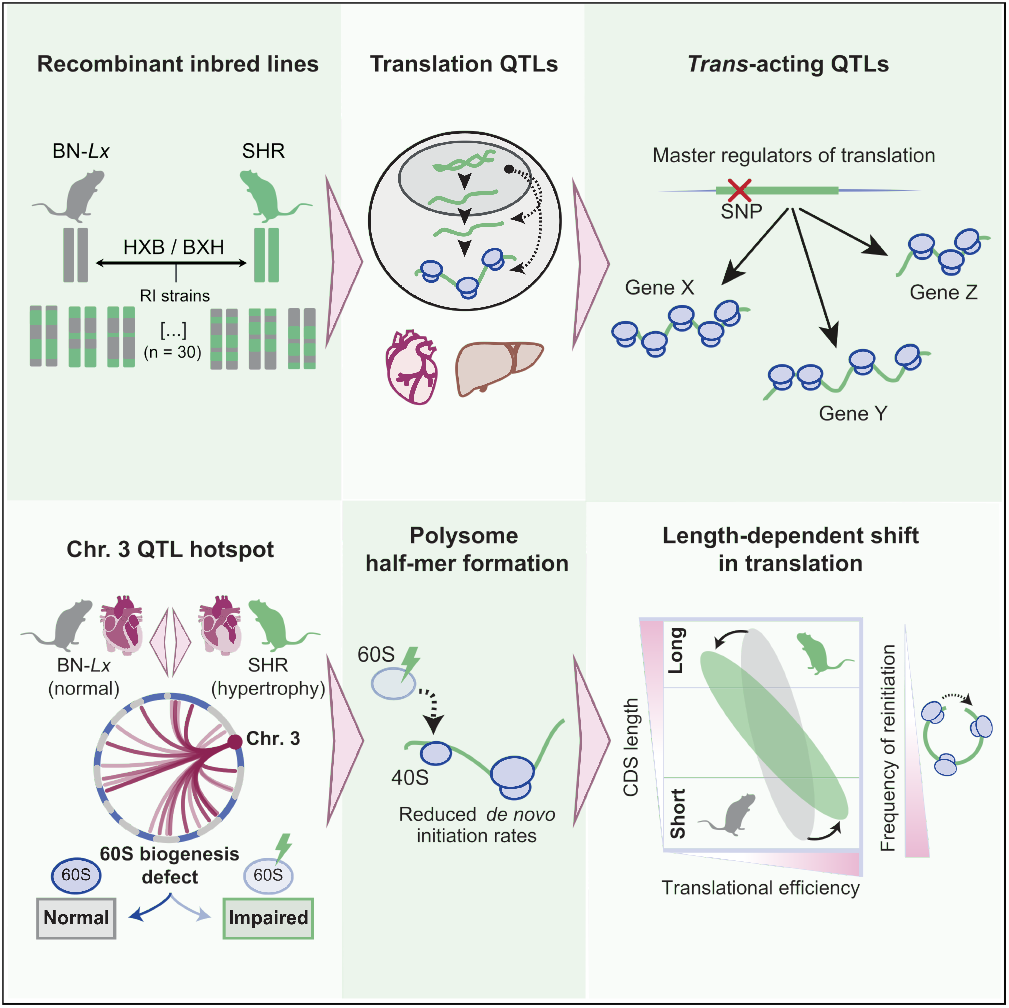

**Highlights:**

- Genetic variability impacts protein synthesis rates in a rat model for cardiac hypertrophy
- A *trans* locus affects stoichiometric translation rates of cardiac sarcomeric proteins
- This master regulator locus induces a global, protein length-dependent shift in translation
- Dysregulated ribosome assembly induces half-mer formation and affects translation initiation rate

## INTRODUCTION

Gene expression regulation is a multilayered process and variation at any level can influence susceptibility to disease (Albert & Kruglyak, 2015; Ward & Kellis, 2012). Heritable, naturally occurring genetic variation can induce gene expression changes through epigenetic (Kasowski et al., 2013; McVicker et al., 2013; Rintisch et al., 2014), transcriptional (Brem et al., 2002; GTEx Consortium et al., 2017; Hubner et al., 2005) and translational (Albert et al., 2014; Battle et al., 2015; Cenik et al., 2015; Muzzey et al., 2014) mechanisms. However, the extent to which *trans*-acting factors influence mRNA translation and thereby contribute to phenotypic diversity between individuals, and possibly complex disease, is not known. In this study, we use the rat HXB/BXH recombinant inbred (RI) panel to identify distant genetic effects on mRNA translation in a complex disease relevant setting. The HXB/BXH panel is a powerful model system for rat genetics that consists of 30 RI lines derived from crossing normotensive Brown Norway (BN-*Lx*) and spontaneously hypertensive rats (SHR/Ola; hereafter SHR) (Printz et al., 2003). Each of these 30 RI lines possesses a homozygous mixture of the ± 3.6 million genetic positions that discriminate both parental lines - an extent of genetic variability resembling that of two unrelated human individuals (Hermsen et al., 2015; Simonis et al., 2012). Within the HXB/BXH panel, these genetic variants can be associated with physiological and molecular phenotypes to uncover disease-relevant genotype-phenotype relationships (Heinig et al., 2010; McDermott-Roe et al., 2011; Petretto et al., 2008; Pravenec et al., 2008). Importantly, for each of the two parental genotypes (BN-*Lx* and SHR), any genetic locus is on average replicated by 15 out of 30 RI lines, providing sufficient power to detect not only local (*cis*) but also distant, *trans*-acting QTLs.

Here we define the influence of genetic variation on the efficiency of mRNA translation (translational efficiency, or TE) by applying ribosome profiling (or Ribo-seq (Ingolia et al., 2009)) and RNA-seq to liver and left-ventricular heart tissue of each of the 30 RI lines - two tissues directly related to the cardiovascular and metabolic traits present in SHR. Focusing specifically on distant translational efficiency QTLs (teQTLs), we discover a prominent set of *trans*-acting ‘hotspots’ that each control the translation of up to dozens of genes in the rat heart. Amongst these potential translational master regulators we find a single distant teQTL on rat chromosome 3 that influences TE in a translatome-wide and protein length-dependent fashion. In-depth investigation of this locus, which colocalizes with a highly replicated locus for left ventricular mass (Inomata et al., 2005; McDermott-Roe et al., 2011; Siegel et al., 2003), reveals a likely defect in ribosome biogenesis that induces half-mer formation indicative of impaired translation initiation, and misregulates the highly abundant small nucleolar RNA *SNORA48*. This ribosomopathy is specifically observed in SHR hearts and reinforces a protein length-dependent imbalance in protein synthesis rates that exists at baseline (Arava et al., 2003; Arava et al., 2005; Ciandrini et al., 2013; Rogers et al., 2017; Shah et al., 2013), but is amplified in disease.

With our work, we show how cardiac translation is under extensive distant genetic control by a limited number of master regulatory loci. We highlight a single genetic locus that induces a complex, translation-driven molecular mechanism that contributes to phenotypic diversity and underlies a complex cardiac trait.

## RESULTS

### Identification of translational efficiency QTLs in the HXB/BXH panel

To be able to associate genetic variation with translational efficiencies, we further refined a previously constructed (STAR Consortium et al., 2008; Rintisch et al., 2014) genotype map of the HXB/BXH RI panel (see **Methods** and **Figure 1A**). The obtained genotypes were associated with the mRNA expression and translation levels of 10,531 cardiac and 9,336 liver genes (77% overlap), which were obtained using Ribo-seq and RNA-seq data across each of the 30 RI lines (**Figure 1B-C, Figure S1A-H** and **Table S1**). We identified and categorized three types of QTLs per tissue: mRNA expression QTLs (eQTLs; mRNA-seq levels), ribosomal occupancy QTLs (riboQTLs; Ribo-seq levels) and translational efficiency QTLs (teQTLs; Ribo-seq levels corrected for mRNA-seq levels) (**Figure 1D-E**, and **Table S2**). In line with previous work (Albert et al., 2014; Battle et al., 2015; Cenik et al., 2015; Heesch et al., 2019), we found that most local QTLs had a clear transcriptional basis (i.e. as eQTLs) that was, with minor variations, concordantly visible in the Ribo-seq data (**Figure S1I**). However, *cis* effects specific to translation (local teQTLs) did exist and clearly influenced a subset of genes (n_heart_ = 71 and n_liver_= 88), driving expression changes independent of mRNA expression regulation (**Figure S1I** and **S1J**). These teQTLs showed limited recurrence between heart and liver, despite most genes with teQTLs being expressed in both tissues (see **Methods**, **Figure 1F** and **Table S2**).

**Figure 1:**
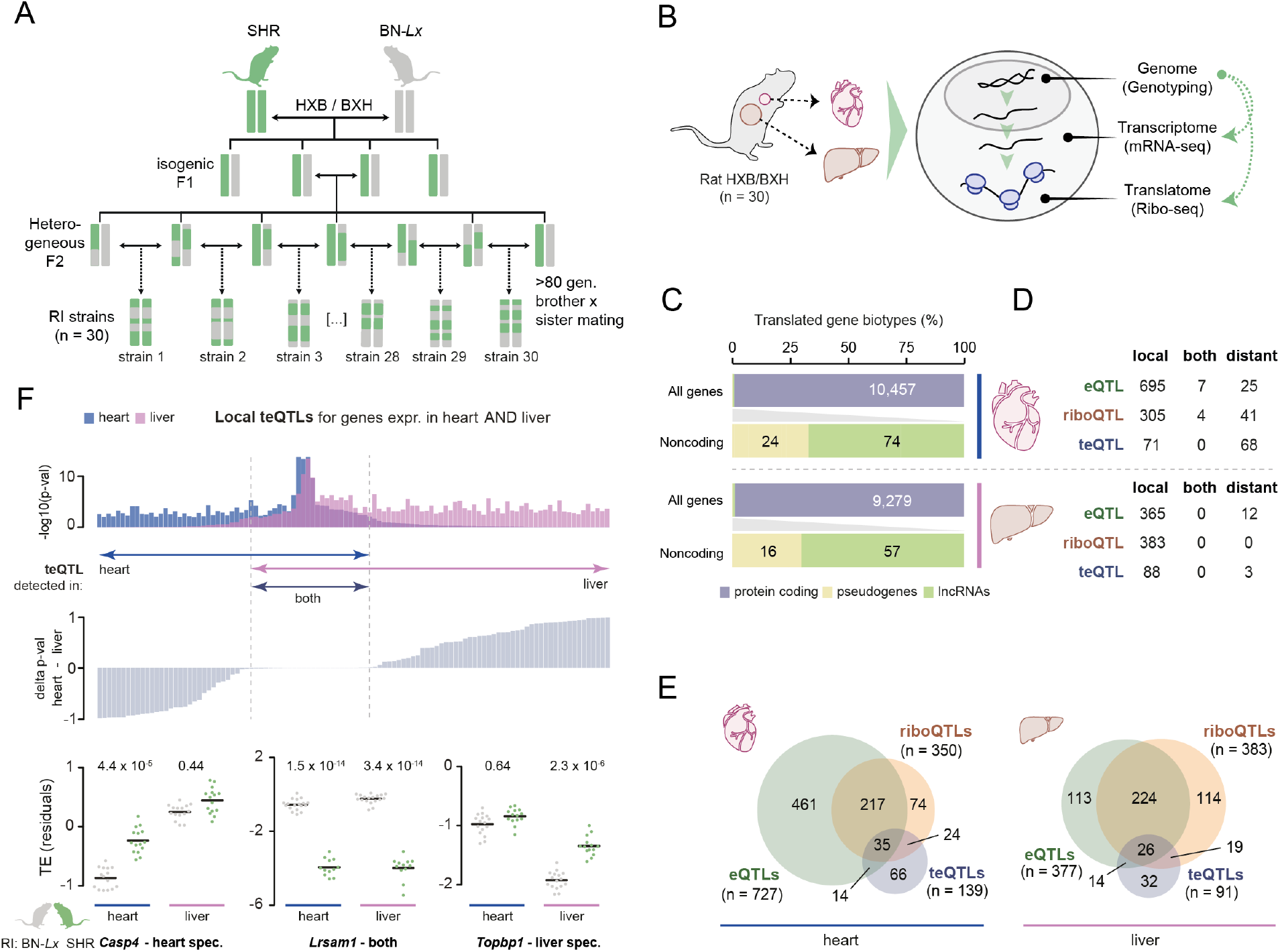
Identification of translational efficiency QTLs in the HXB/BXH panel. **(A)** Schematic overview of the establishment of the HXB/BXH recombinant inbred panel. Colored bars represent SHR and BN-*Lx* alleles. **(B)** Schematic overview of the experimental procedures carried out for each of the 30 HXB/BXH lines. **(C)** Bar plot illustrating absolute and relative ORF identifications, separated by coding and noncoding gene biotypes, for heart and liver. Translated pseudogenes were excluded from downstream analyses. See also **Table S1**. **(D)** Table with gene-centric QTL mapping results for heart and liver, separated by genes for which mRNA expression QTLs (eQTLs), ribosome occupancy QTLs (riboQTLs), and/or translational efficiency QTLs (teQTLs) are identified. Local QTLs indicate associations that map to the same genomic locus as the tested gene (see **Methods**). Distant QTLs refer to associations with genes on chromosomes other than that of the QTL. See also **Table S2**. **(E)** Venn diagrams displaying a gene-centric overlap of eQTLs, riboQTLs and teQTLs in heart and liver, highlighting QTLs shared with, or specific to, a single trait. **(F)** Bar plot with a tissue comparison of detected local translational efficiency QTLs (teQTLs) considering only genes expressed and translated in both tissues. Genes are ordered by the delta p-value in heart vs. liver tissue (middle panel). Three examples of genes expressed in heart and liver tissue are given, displaying a local teQTL in either one, or both, of these tissues. Cross bars indicate mean values. See also **Figure S1** and **Tables S1-S3**.

While this is possibly explained by liver being a frequent outlier in cross-tissue eQTL comparisons (Aguet et al., 2019), these findings suggest that many *cis*-acting teQTLs are mediated in a tissue-specific manner.

### Local teQTLs are mechanistically independent of upstream ORFs

Upstream ORFs (uORFs) are major regulatory elements of translation located in 5’ leader sequences of protein-coding mRNAs (Morris & Geballe, 2000), and genetic variants interfering with these elements can affect the efficiency of mRNA translation (Cenik et al., 2015). Out of over a thousand newly detected uORFs per tissue (**Figure S1H** and **Table S3**), we detected 27 (heart) and 13 (liver) uORFs whose translation rates associated with genetic variants in *cis* (“uORF-QTLs”; **Table S3**). However, none of these variants disrupted the uORF’s start or stop codon, and only a single uORF-QTL colocalized with a primary ORF teQTL. For this gene, *Rte1*, both QTLs showed the same effect directionality, indicating that increased translation of the uORF had no negative impact on the primary ORF TE (**Figure S1K**). In general, uORF and primary ORF translation rates showed a very limited quantitative dependency (as observed in (Aspden et al., 2014; Brar et al., 2012; Chew et al., 2016; Heesch et al., 2019)) (**Table S3**, **Figure S1L** and **S1M**) and we found no enrichment of uORFs in genes with local teQTLs (p_heart_ = 0.70 and p_liver_ = 0.79). In addition, we found no genetic variants in genes with local teQTLs that interfered with local translation initiation context or Kozak sequence. Although we cannot determine the possible outcome of genetic variants in other functional elements that serve to fine-tune mRNA translation, such as RNA folding structures, methylation sites, or RNA binding protein motifs (Hershey et al., 2012; Wang et al., 2015), our observations imply that uORFs are unlikely to be main drivers of local teQTLs within the HXB/BXH panel.

### Distant teQTL “hotspots” are master regulators of cardiac translation

Distant QTLs are an important source of variation in mRNA expression levels, through which they contribute to complex disease (Aguet et al., 2019; Brandt et al., 2020).

Although the impact of *trans*-acting QTLs on mRNA translation in a complex disease setting has remained unexplored, the HXB/BXH panel employed here provides enough power to detect such QTLs (Aitman et al., 1999; Heinig et al., 2010; Hubner et al., 2005; Pravenec et al., 2008). Because we found distant teQTLs to be more frequent in heart than in liver (**Figure 1D**), we decided to focus downstream analyses solely on heart tissue. To increase the power to detect genes with shared modes of regulation by a single QTL "hotspot", we applied a hierarchical regression model in a Bayesian framework using a stochastic search algorithm (HESS (Bottolo et al., 2011b; Lewin et al., 2016)) (see **Methods**). This yielded a higher total of 243 genes whose TE is regulated by distant teQTLs (**Figure S2A** and **Table S4**). Of all distant teQTLs, we classified nine loci as distant cardiac master regulators, as they influenced the TE of at least 5 (but up to 25) genes distributed over different chromosomes (**Figure 2A**, **Figure S2B** and **Table S4**).

**Figure 2:**
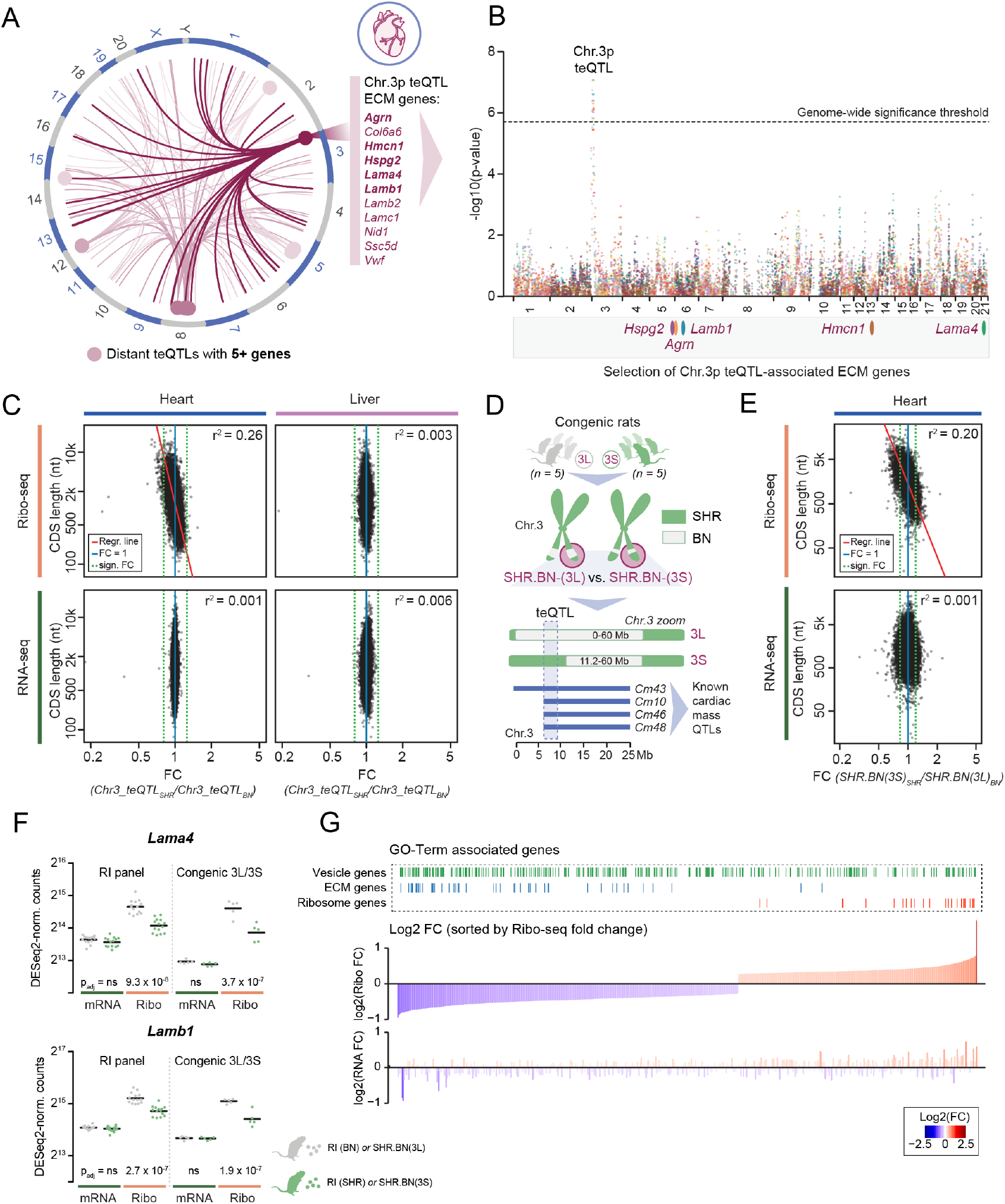
The chromosome 3p teQTL regulates cardiac translation in a protein length-dependent manner. **(A)** Circos plot highlighting all distant teQTLs detected in the rat heart that associate with the TE of at least 5 genes. The Chr. 3p teQTL is highlighted in dark pink and of the 25 associated genes, only the names of the 11 extracellular matrix (ECM) genes are given. Gene-locus associations for the different hotspots are indicated with the different shades of pink, with the darkest pink and thickest lines representing the associations identified for the Chr. 3p teQTL. For each hotspot, translational efficiencies of all associated genes are given in **Figure S2B**. **(B)** Overlay of Manhattan plots displaying genome-wide significance values for a genetic association with TE on Chr. 3p. A selection of 5 associated genes whose protein products function in the extracellular matrix are shown. **(C)** Scatter plots and square correlation coefficients (r^2^) based on standardized major axis (SMA) values between coding sequence (CDS) length and the fold change (FC) in gene expression, as measured by Ribo-seq (top) or mRNA-seq (bottom), for heart (left) and liver tissue (right). To define the expression FC, all 30 RI lines are separated by local genotype (BN or SHR) at the Chr. 3p teQTL. For heart Ribo-seq data, the correlation is significant (p-value < 2.2 10^−16^; Test of correlation coefficient against zero) and the linear model based on fitted SMA method is displayed as a red line. **(D)** Schematic overview of the congenic rat lines with isolated teQTL and cardiac mass QTL locus. The SHR.BN-(3L) line carries a local BN genotype, whereas the SHR.BN-(3S) line retains the SHR genotype at the teQTL. Inserted BN segments are visualized in grey, SHR alleles in green. **(E)** Scatter plots and square correlation coefficients (r^2^) based on standardized major axis (SMA) values between coding sequence (CDS) length and the fold change (FC) in gene expression, as measured by Ribo-seq (top) or mRNA-seq (bottom) in congenic rat hearts. The FC in translation is derived from a comparison between 5 replicates of SHR.BN-(3L) and SHR.BN-(3S) rats and reproduces the global length effect observed for the Chr. 3p teQTL identified in the HXB/BXH RI panel. For heart Ribo-seq data, the correlation is significant (p-value < 2.2 10^−16^; Test of correlation coefficient against zero) and the linear model based on fitted SMA method is displayed as a red line. **(F)** Dot plots with indications of mean expression for 2 laminin subunits (extracellular matrix glycoproteins), illustrating the reproducibility of the translational efficiency phenotype between the HXB/BXH RI panel and the congenic rat lines. Cross bars indicate mean values. **(G)** Bar plot with all differentially translated genes in a comparison of both congenic rat lines, ordered by Ribo-seq FC in expression. Genes associated with selected significant GO terms are highlighted on top. See also **Figure S2** and **Table S4**.

A single 2.9 Mb large teQTL hotspot on rat chromosome 3p (Chr. 3: 6.3 - 9.2 Mb; equivalent to human Chr. 9q34) drew our attention for being associated with the TE of 25 genes (**Figure 2A**, **Figure S2B**, **Table S2** and **S4**). This locus furthermore co-localized with a highly replicated QTL for cardiac mass (Inomata et al., 2005; Siegel et al., 2003), for which a loss-of-function insertion in endonuclease G (*Endog*) was previously identified as a driver of cardiomyocyte hypertrophy and increased left-ventricular weight (McDermott-Roe et al., 2011). Among all genes associated with this master regulatory teQTL, we found strong enrichment for extracellular matrix (ECM) proteins (11 out of 25; GO:0031012; p_adj_ = 3.32 × 10^−10^) (**Figure 2A** and **2B**), consistent with recent observations of strong translational control of fibrotic processes in human hearts (Chothani et al., 2019; Heesch et al., 2019).

### The chromosome 3p teQTL regulates cardiac translation in a protein length-dependent manner

The strong translational impact on ECM genes led us to hypothesize that the differential translation could be related to a global switch in translational control related to the generally high coding-sequence (CDS) length of ECM proteins. Indeed, we observed a moderate, though significant correlation between CDS length and fold change (FC) in translation (r^2^ = 0.26), which produces a downregulatory effect for genes with long CDSs and, vice versa, an upregulatory effect for genes with short CDSs (**Figure 2C**). This association with CDS length was specific to heart tissue, absent in RNA-seq data, and no other genetic locus outside of the Chr. 3p teQTL showed a similar effect.

To replicate this translatome-wide phenotype, we performed ribosome profiling on two congenic rat lines with two small, but differently sized, BN segments inserted into the short arm of Chr. 3 on an otherwise fully SHR background (McDermott-Roe et al., 2011) (see **Methods** and **Figure 2D**). The first congenic line possessed a long BN segment that replaced the teQTL completely (SHR.BN-(3L)), whereas the second line contained a smaller BN segment positioned adjacent to the teQTL (SHR.BN-(3S)), hence leaving the teQTL intact. Comparing the cardiac translatomes of both congenic lines, we fully recapitulated the protein length-dependent difference in translation observed in the HXB/BXH RI panel (r^2^ = 0.20; **Figure 2E+F**). A subsequent GO enrichment analysis on differentially translated genes concordantly yielded terms matching the downregulation of very large proteins (GO: extracellular region; p_adj_ = 6.33 × 10^−13^) or the upregulation of very small proteins (GO: cytosolic ribosome; p_adj_ = 1.22 × 10 ^−13^) (**Figure 2G**). Of note, the observed TE fold changes specifically correlated with CDS length (r^2^ = 0.20), to a lesser extent with total transcript length (r^2^ = 0.162) but not with 5’ UTR (r^2^ = 0.004) or 3’ UTR length (r^2^ = 0.013) (**Figure S2C**).

### The chromosome 3p teQTL induces polysome half-mer formation

To mechanistically dissect the translational phenotype linked to the Chr. 3p teQTL, we next performed polysome profiling on heart tissue of both congenic lines (**Figure 3A**). Polysome profiles of SHR.BN-(3S) rats showed heavily altered ribosomal configurations compared to SHR.BN-(3L) (**Figure 3A+B** and **Figure S3A**), exemplified by “shoulders” accompanying each polysome peak indicative of polysome half-mer formation (**Figure 3C**). Polysome half-mers are formed when the 43S preinitiation complex does not instantly join the large 60S ribosomal subunit to form a functional 80S monosome. This stalls translation initiation - the rate-limiting step of RNA translation and therefore a main determinant of TE (Gandin et al., 2008; Shah et al., 2013; Sonenberg & Hinnebusch, 2009). Half-mers arise because of ribosome biogenesis defects, caused by the underproduction of 60S subunits (Rotenberg et al., 1988) or impaired subunit joining (Colón-Ramos et al., 2006; Eisinger et al., 1997). However, for all investigated rats ribosomal subunit production levels appeared balanced (**Figure S3B**).

Globally remodeled translatomes can also result from proteotoxic stress at the endoplasmic reticulum (ER). Although ER stress and the unfolded protein response play a pivotal role in the pathophysiology of the heart (Groenendyk et al., 2010; McLendon & Robbins, 2015), we found no characteristic signatures of such a response: the polysome profiles did not resemble those typically observed upon ER stress (a strong shift from polysomal to predominantly monosomal translation (Baird et al., 2014; Harding et al., 2000)) (**Figure 3A**), translation elongation rates remained constant along the entire CDS (Liu et al., 2013) (**Figure S3C**), protein levels of common ER stress markers (e.g. p-IRE1*α*:IRE1*α* ratios and XBP1s (Schiattarella et al., 2019)) were unchanged **(Figure S3D**), mRNA expression and translation levels of genes associated to ER stress (GO: 0034976) or apoptosis (GO: 0097190) were not affected (p = 0.70 and 0.32, respectively), and translationally upregulated genes were not enriched for having uORFs (Guan et al., 2017) (**Figure S3E**).

**Figure 3:**
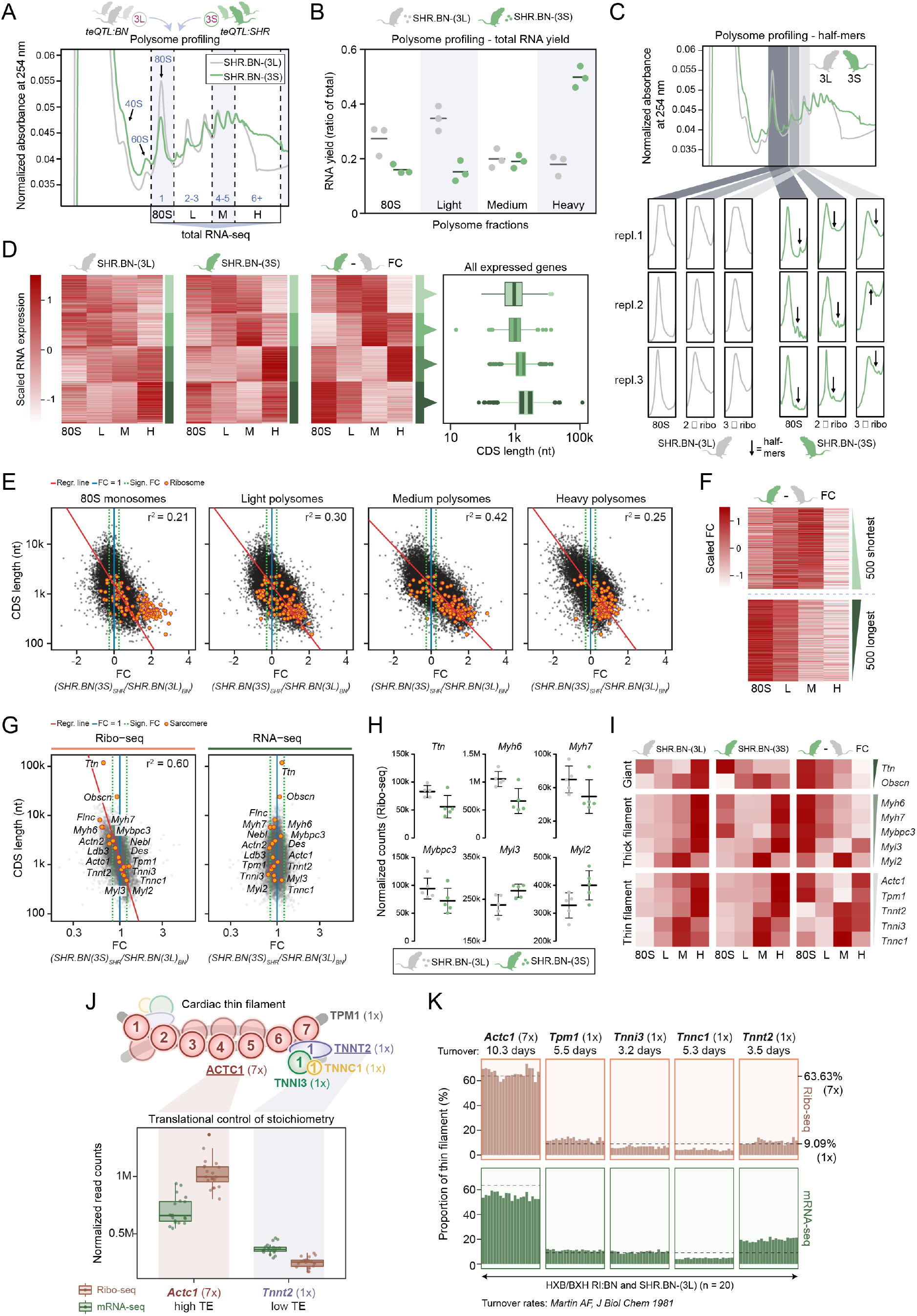
The chromosome 3p teQTL induces polysome half-mer formation. **(A)** Schematic overview of the polysome fractionation and RNA-seq approach. One representative polysome profile per congenic rat line is given. L, M and H fractions indicate light, medium and heavy polysomes, respectively. **(B)** Congenic line comparison for differences in polysomal configuration as measured by the distribution of RNA yield across the fractions. Quantified polysome profile area under curves (AUCs) can be found in **Figure S3A**. Bars indicate mean values. **(C)** Zoomed-in view of multiple polysomal peaks across replicates for both congenic lines, with arrows indicating half-mers. **(D)** Heatmap with scaled RNA-seq expression levels of all 12,471 quantified genes (mean RNA FPKM ≥ 1 across replicates, for both lines). Genes are clustered into 4 groups by k-means clustering, and sorted by CDS length within each cluster. The same gene order obtained through clustering of the fold-change (SHR.BN-(3S) vs SHR.BN-(3L)) comparison (3^rd^ heatmap) was used for the individual heatmaps of SHR.BN-(3L) vs SHR.BN-(3S) (1^st^ and 2^nd^ heatmap). For all clusters, box plots with the CDS length distribution are shown on the right. **(E)** Scatter plots and square correlation coefficients (r^2^) based on standardized major axis (SMA) values between coding sequence (CDS) length and the fold change in gene expression (FC (SHR.BN-(3S) vs SHR.BN-(3L)), as measured by RNA-seq of the four isolated fractions. The correlations are significant (p-value < 2.2 × 10^−16^; Test of correlation coefficient against zero) and the linear model based on fitted SMA method are displayed as red lines. Ribosomal protein genes (with small CDSs) are depicted by orange dots. **(F)** Heatmaps with the scaled FC of the ribosomal configuration of the top 500 shortest and longest CDS genes. **(G)** Scatterplots showing CDS length versus fold change (FC (SHR.BN-(3S) vs SHR.BN-(3L)) for Ribo-seq and RNA-seq data, highlighting a representative selection of core- and accessory sarcomere proteins. The square correlation coefficient (r^2^) based on standardized major axis (SMA) is calculated using expression values of this subset of genes only. **(H)** Dot plots with Ribo-seq expression values for *Ttn* and a selection of cardiac thick filament proteins. Genes are sorted by CDS length from top left to bottom right. Error bars indicate mean values with standard deviation (SD). None of the displayed expression changes are genome-wide significant. **(I)** Heatmaps with polysome profiling results for selected sarcomere proteins. Expression distributions for the individual animals, as well as the scaled fold changes between SHR.BN-(3S) vs SHR.BN-(3L) are given. Within each group, genes are sorted by CDS length (top to bottom). **(J)** Schematic representation of the cardiac thin filament and its composition stoichiometry as obtained from (Thompson & Metzger, 2014). Cardiac muscle alpha actin (*Actc1*) and cardiac troponin T (*Tnnt2*) are the genes most strongly translationally regulated to achieve desired protein levels. **(K)** Bar plots showing the relative contribution of each thin filament component as measured by Ribo-seq (top) and mRNA-seq (bottom) expression levels. DESeq2-normalized expression values are corrected for reported rat heart protein turnover rates (Martin, 1981) and represented as a percentage of the complete thin filament. Twenty healthy rats are shown (from left to right: 5x SHR.BN-(3L) congenic animals, followed by 15x HXB/BXH RI lines as separated by local BN genotype according to the Chr. 3p teQTL). Optimal production values for 7 or 1 subunit(s) are indicated by dashed lines. See also **Figure S3**.

### The chromosome 3p teQTL influences stoichiometric sarcomere translation

Comparing RNA-seq data from isolated fractions of monosomes (80S), light-(2-3 ribosomes), medium-(4-5 ribosomes) and heavy-weight polysomes (6+ ribosomes), we again saw a clear relationship between ribosome occupancy and CDS length (**Figure 3D**). This length-dependency was identical to the one observed in the Ribo-seq data, validating the TE phenotype through an independent method (**Figure 3E**). Whereas mRNAs with the longest CDSs showed a clear reduction in heavy polysome occupancy, accompanied by a relative enrichment in the monosomal fraction, mRNAs with the shortest CDSs showed increased steady-state translation in light- and medium polysomal configurations (**Figure 3F**).

Among the genes most strongly affected by the length-dependent shift in ribosomal occupancy and TE were multiple core sarcomere proteins (**Figure 3G-I**). These primarily included ‘giant’ proteins *Ttn* and *Obscn*, as well as the larger protein constituents of the thick (*Myh6*, *Myh7* and *Mybpc3*) and thin filament (*Actc1* and *Tpm1*), which all showed downregulated translation. In contrast, the much smaller components of the thick and thin filament, such as the myosin light chains (*Myl2* and *Myl3*) and cardiac troponins (*Tnnc1*, *Tnnt2* and *Tnni3*), were all translationally upregulated. The large variability in sarcomere protein sizes correlated well with translational fold change (r^2^_sarcomere_ = 0.60; **Figure 3G**), highlighting the impact of the Chr. 3p teQTL on sarcomere gene translation.

Of note, sarcomere homeostasis strongly depends on stoichiometric protein production and mRNA translation has been proposed to regulate this equilibrium (Palermo et al., 1996; Rethinasamy et al., 1998). For the cardiac thin filament in particular, we indeed saw prominent translational control of protein production, exemplified by the translational up- and downregulation of *Actc1* (TE = 1.50) and *Tnnt2* (TE = 0.69), respectively, to achieve protein production levels in compliance with composition stoichiometry (**Figure 3J+K**). Pushing the normally proportional filament translation rates into opposite directions because of differences in subunit CDS lengths (**Figure 3G-I**), it becomes challenging to achieve composition stoichiometry in an energy efficient manner (Li et al., 2014; Taggart & Li, 2018), as such imbalances need to be corrected post-translationally through the targeted degradation of excess subunits (McShane et al., 2016; Taggart et al., 2020).

### Reduced *de novo* translation rates reinforce a pre-existing length-bias in TE

Having established that the Chr. 3p teQTL influences TE through changes in ribosomal configurations and the formation of polysome half-mers, it remains unclear why the severity of this phenotype correlates with protein length. It is known that the density of ribosomes along mRNAs shows a translatome-wide inverse correlation with CDS length and, as a consequence, longer proteins are generally less efficiently translated than shorter ones (Arava et al., 2003; Arava et al., 2005; Ciandrini et al., 2013; Rogers et al., 2017). This length-effect is directly linked to the frequency of translation reinitiation, which decreases with increasing CDS length (Ciandrini et al., 2013; Rogers et al., 2017; Shah et al., 2013). Indeed, in both unaffected SHR.BN-(3L) and SHR.BN-(3S) rat hearts, TE correlates negatively with CDS length (r^2^ = 0.12 and r^2^ = 0.21 respectively; **Figure 4A**). Upon limited or hampered *de novo* assembly of 80S monosomes, SHR.BN-(3S) rats become increasingly dependent on effective ribosome recycling (Rogers et al., 2017), for which both previously acquired ribosomal subunits remain instantly available. In agreement with this, the effect on TE is significantly enhanced in SHR.BN-(3S) rats (Fisher r-to-z transformation Z = −11; p < 2.2 10^−16^) (**Figure 4A**). This effect is less detrimental for mRNAs with short CDSs, which more frequently reinitiate as one round of translation takes less time to complete, thereby reinforcing a pre-existing length-dependency in translation (**Figure 4B**).

**Figure 4:**
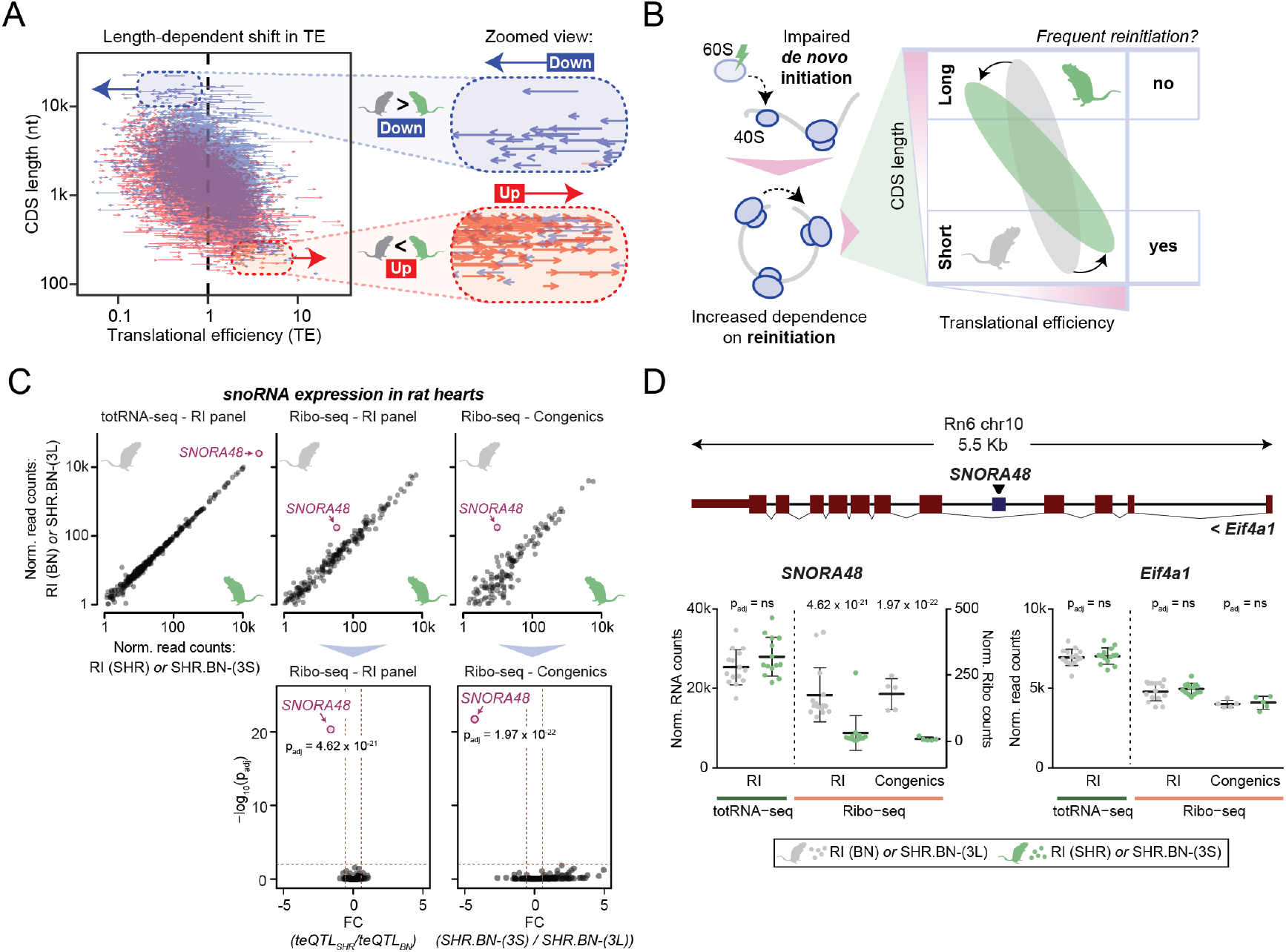
Reduced *de novo* translation rates reinforce a pre-existing length-bias in TE. **(A)** Arrow-based scatter plot show the transitions in TE per gene, between SHR.BN-(3S) and SHR.BN-(3L) rats. The length of the arrow is representative of the absolute change in TE between both congenic lines, with the position of the arrow tail reflecting the SHR.BN-(3L) TE and the position of the arrow head indicating the TE in SHR.BN-(3S) rats. Blue arrows indicate a decrease in TE in SHR.BN-(3S) rats, whereas red arrows indicate an increase in TE in SHR.BN-(3S). Two zoomed-in regions show arrow behavior in the top and bottom of the graph, respectively highlighting genes with very long and short CDSs. **(B)** Schematic of how the ribosome biogenesis defect leads to a change in translation initiation rates and a subsequent global shift in TE that correlates with CDS length. **(C)** Scatter plots showing expression levels of all cardiac-expressed snoRNAs as measured by totRNA-seq and Ribo-seq data, with *SNORA48* highlighted in pink. For both Ribo-seq datasets, p-value volcano plots show the significance of the differential regulation of *SNORA48* (highlighted in pink). **(D)** Representation of the genomic location of *SNORA48*. This snoRNA is contained within intron 4 of its host gene *Eif4a1*. Dot plots with expression levels as measured by totRNA-seq and Ribo-seq for *SNORA48* and its host gene *Eif4a1*, in both the HXB/BXH RI panel and the congenic rat lines. Error bars indicate mean values with standard deviation (SD). See also **Figure S4**.

### Misregulation of pseudouridylation guide SNORA48 characterizes the Chr. 3p teQTL

We next searched for possible misregulated ribosome biogenesis factors that could explain the observed phenotype, since mutations in these elements are known to induce global changes in polysome profiles similar to what we observe in the affected hearts (Li et al., 2009; Tafforeau et al., 2013).

We find one such factor differentially regulated in the affected hearts: the H/ACA box small nucleolar RNA (snoRNA) *SNORA48* (also known as *ACA48*; Ensembl ID ENSRNOG00000060816). *SNORA48* is a conserved snoRNA predicted to guide the pseudouridylation (Ψ) of 28S rRNA during large ribosomal subunit biogenesis (Lestrade & Weber, 2006) (**Figure S4A**). It is the most highly expressed snoRNA in rat hearts (**Figure 4C**) and the only snoRNA that shows a genome-wide significant decrease in ribosome association, while overall production levels of the snoRNA and the host gene *Eif4a1* remain constant (**Figure 4D**).

To identify the cause of *SNORA48* misregulation, we investigated the involvement of the key Chr. 3p teQTL candidate gene *Endog*, whose loss-of-function (LoF) increases cardiac mass and impairs cardiac energy metabolism (McDermott-Roe et al., 2011) (**Figure S4B**). We profiled the cardiac translatomes of wild type and knockout *Endog* mice (obtained from (McDermott-Roe et al., 2011)), as well as those of wild type and newly established transgenically rescued Endog SHR rats (**Table S1** and **Methods**).

Unfortunately, these models showed no reduced ribosomal association of *SNORA48* and no clear length-dependent translational phenotype (**Figure S4C**). This excludes Endog as a monogenic driver of the Chr. 3p teQTL and points to other mutated genes in the locus. Prime locus candidates with predicted damaging mutations include the DEAD box helicase *Ddx31* (yeast *DBP7*), whose deletion reduces 60S levels and induces half-mer formation in yeast (Daugeron & Linder, 1998; Tafforeau et al., 2013), the ribosomal RNA transcription termination factor *Ttf1* (Grummt et al., 1985), or the methyltransferase *Spout1* (also known as *C9ORF114*), which codes for an essential pre-rRNA processing factor (Tafforeau et al., 2013). Although our data indicate that *Endog* does not act autonomously in the establishment of the Chr. 3p teQTL, complex genetic interactions with one or more of these dysfunctional ribosome biogenesis genes may be required for the translational phenotype to arise.

## DISCUSSION

In this study, we use a QTL mapping strategy to define the influence of natural genetic variation on the efficiency of mRNA translation, with a focus on identifying distant, *trans*-acting QTLs that control multiple genes. Genetic influences on translation have previously been studied in yeast (Albert et al., 2014; Muzzey et al., 2014) and a cohort of human lymphoblastoid cell lines (LCLs) (Battle et al., 2015; Cenik et al., 2015), albeit solely carried out in in vitro culture systems with limited focus on distant QTLs. These studies show that within the investigated systems, the vast majority of (local) eQTLs are fed forward into variation in protein levels, with limited specific impact on translation ((Albert et al., 2014; Battle et al., 2015; Cenik et al., 2015; Muzzey et al., 2014); reviewed in (Albert & Kruglyak, 2015; Liu et al., 2016)). Although we similarly see high concordance between local genetic influences on mRNA expression and translation, we do detect multiple teQTLs with specific and prominent effects on the mammalian tissue gene expression landscape. The most apparent effects are orchestrated through a limited set of distant master regulatory loci - or teQTL “hotspots” each controlling the TE of up to dozens of genes. This widespread distant translational control is particularly abundant in the heart and likely crucial for the adaptation to developing (patho)physiological conditions, though may be dormant (and hence go undetected) in unaffected tissue.

We mechanistically dissect a prominent distant teQTL on rat chromosome 3 that drives a translatome-wide and protein length-dependent change in TE. We show that this teQTL induces a global shift in ribosome configurations and triggers the formation of polysome half-mers, which we trace back to a likely ribosome biogenesis defect that impairs *de novo* translation initiation. To understand the basis of this specific molecular phenotype and the consequences for heart disease, it is important to know that length-dependent differences in the efficiency of translation are present at baseline in the translatomes of all species (Ciandrini et al., 2013; Rogers et al., 2017; Shah et al., 2013). This phenomenon has been directly connected to the rate of translation initiation (Ciandrini et al., 2013; Rogers et al., 2017; Shah et al., 2013) and is most likely explained by varying rates of translation reinitiation (Rogers et al., 2017). As a single round of translation at a short CDS takes less time to complete, reinitiation rates are higher, which ultimately yields more protein. Hence, when *de novo* initiation rates are reduced because of a ribosome biogenesis defect, this does not necessarily decrease the efficiency of translation reinitiation, as both subunits have already been recruited and properly assembled once. It does make mRNAs more dependent on effective and frequent reinitiation for their translational output, thereby enhancing a pre-existing length-dependent imbalance in TE which is exactly what we observe in the rat hearts that carry the SHR genotype at the Chr. 3p teQTL (**Figure 4A+B**).

Interestingly, deletion of *SNORD24* was previously shown to induce a half-mer (Kouba et al., 2012; Li et al., 2009) and length-specific translational phenotype (Thompson et al., 2016) very similar to what we observe in this study. *SNORD24*, also known as *SNR24* or *U24*, is a highly conserved C/D box snoRNA required for the site-specific 2’-O-methylation of 28S rRNA during 60S ribosomal subunit biogenesis (Kiss-László et al., 1996; Kouba et al., 2012; Li et al., 2009; Qu et al., 1995). Its deletion in yeast reduces 60S levels and subsequently induces a polysome half-mer phenotype accompanied by a translatome-wide and length-dependent shift in TE (Kouba et al., 2012; Li et al., 2009; Thompson et al., 2016). In line with these results, one of the most striking gene expression changes we observe in association with the Chr. 3p teQTL is the reduced ribosomal association of *SNORA48*. Similar to *SNORD24*, *SNORA48* is predicted to guide the modification of 28S rRNA and its high abundance in the rat heart could indicate that its misregulation is equally detrimental for 60S ribosomal subunit biogenesis. The high similarity between both phenotypes raises the possibility that length-dependent changes in TE more commonly result from ribosome biogenesis defects that induce polysome half-mer formation. Even though all ribosomopathies originate from defects in ribosome biogenesis, they often lead to unique phenotypes with tissue-specific clinical manifestations (Danilova & Gazda, 2015). This potential and thus far overlooked consequence of translational deficiencies shows apparent conservation from yeast to mammals, and could be an important mediator of the molecular changes that connect common ribosomopathies with specific clinical symptoms (Narla & Ebert, 2010).

Our work shows that naturally occurring genetic variation can induce a complex, translation-driven molecular mechanism that globally reforms mammalian tissue gene expression. Distant genetic control of mRNA translation is frequent and contributes significantly to interindividual phenotypic variability, coordinated by multiple master regulatory loci that each regulate the TE of multiple genes. We anticipate that adaptation of gene expression regulation through mRNA translation is crucial for tissues developing complex phenotypic traits and highlight a single locus that influences TE in a protein length-dependent fashion, likely as a result of a ribosome biogenesis defect that affects translation initiation.

## Author Contributions

Conceptualization, S.v.H. and N.H.; Methodology, S.v.H.; Software, F.W. and J.R.O; Validation, J.F.S.; Formal Analysis, F.W., J.R.O., V.S.L., G.P., O.H., L.B., M.V. and M.H.; Investigation, S.v.H., C.C.M., S.B., E.A., M.B.M. and J.F.S.; Resources, J.S., D.S., M.C., and M.P.; Data Curation, F.W., J.R.O., G.P. and O.H.; Writing - Original Draft, S.v.H.; Writing - Review & Editing, S.v.H. and J.R.O., with input from all authors; Visualization, S.v.H., F.W., E.A., C.C.M. and J.R.O.; Supervision, S.v.H.; Project Administration, S.v.H. and N.H.; Funding Acquisition, S.v.H. and N.H.

## Acknowledgements

S.v.H. was supported by an EMBO long-term fellowship (ALTF 186-2015, LTFCOFUND2013, GA-2013-609409). N.H. is the recipient of an ERC advanced grant under the European Union Horizon 2020 Research and Innovation Program (grant agreement AdG788970) and is supported by a grant from the Leducq Foundation (16CVD03). M.P. was supported by Praemium Academiae award (AP1502) of the Czech Academy of Sciences. D.S. was funded by Grant 20153810 from Fundació La Marató de TV3.

## METHODS

### RESOURCE AVAILABILITY

#### Lead contact

Further information and requests for resources and reagents should be directed to and will be fulfilled by the Lead Contact, Norbert Hubner.

#### Materials availability

The transgenic SHR/Ola-Tg(CMV-*Endog*)136 rat line with rescued expression of *Endog* was newly established for this study in the lab of Michal Pravenec (Institute of Physiology of the Czech Academy of Sciences, 142 20, Praha 4, Czech Republic) and is available upon request.

#### Data and code availability

The accession number for the raw rat and mouse sequencing data reported in this paper, normalized and ready-to-use sequencing read count matrices, as well as a further refined genotype map of the HXB/BXH recombinant inbred panel (list of all SDPs and genotypes) is European Nucleotide Archive (ENA): PRJEB38096. All analyses in this study are performed using published and publicly available analytical tools or software packages, with the precise software versions parameters used detailed in the respective **Methods** sections. The code used for the implementation of these tools, as well as all other computational analyses in this study, is available upon request.

### EXPERIMENTAL MODEL AND SUBJECT DETAILS

#### Animal models

Six-week old male HXB/HXB RI rats (n = 30; left ventricle and liver), congenic SHR.BN-D3Rat108/D3Rat56 rats (SHR.BN-(3L); n = 5; left ventricle), congenic SHR.BN-D3Rat108/D3Rat124 rats (SHR.BN-(3S); n = 5; left ventricle), transgenic SHR/Ola-Tg rats expressing *Endog* (n = 5; left ventricle) and wild type SHR/Ola rats carrying an *Endog* LoF mutation (n = 5; left ventricle) were housed, bred and fed ad libitum with a natural diet (Altromin 1314) in an air-conditioned animal facility at the Czech Academy of Sciences, Prague, Czech Republic. All rat experimental procedures were carried out in accordance with the European Union National Guidelines and the Animal Protection Law of the Czech Republic (311/1997) and were approved by the Ethics Committee of the Institute of Physiology, Czech Academy of Sciences, Prague. The congenic rat lines were designed as follows (as described in (McDermott-Roe et al., 2011)): For (SHR.BN-D3Rat108/D3Rat56 or "SHR.BN-(3L)”), a longer genomic BN fragment (Chr3: 0-60 Mb) replaces the entire Chr. 3p teQTL in an otherwise fully SHR/Ola genetic background. For (SHR.BN-D3Rat108/D3Rat124 or "SHR.BN(3S)"), only a shorter fragment (Chr3: 11.2-60 Mb) adjacent to, but not overlapping, the identified teQTL is replaced with a BN fragment. Transgenic SHR/Ola-Tg(CMV-*Endog*)136 strain (hereafter referred to as the SHR-*Endog* transgenic) was derived by microinjecting fertilized eggs with a mix of the Sleeping Beauty construct containing *Endog* cDNA of BN origin under control of the universal EF-1*α* promoter and mRNA of the SB100X transposase. Transgenic rats were detected using PCR with the following primers: *Endog*-F 5’-CGA CAC CTT CTA CCT GAG CA-3’ and *Endog*-R 5’-GGC CCT GTG CAG ACA TAA AC-3’.

The Endog KO mouse was derived from a C57BL/6J background and provided by Dr. Michael Lieber, University of Southern California, LA, CA, USA (Irvine et al., 2005). From the provided founder animals, a colony was established and actively maintained for multiple years within the lab of Daniel Sanchis (Institut de Recerca Biomedica de Lleida, Spain) (McDermott-Roe et al., 2011). The six-week old male mice used for Ribo-seq and RNA-seq experiments were housed in Tecniplast GM500 cages (391 × 199 × 160 mm) never exceeding 5 adults / cage. The investigation with this mouse line was approved by the Experimental Animal Ethic Committee of the University of Lleida (code CEEA N02-02/15) and conforms to the Guide for the Care and Use of Laboratory Animals, 8th Edition, published in 2011 by the US National Institutes of Health.

All rat and mouse animals used in this study were drug and test naive, specific pathogen free (SPF) and not involved in previous procedures.

### METHOD DETAILS

#### Generating a genotype map of the HXB/BXH panel

To refine an existing (STAR Consortium et al., 2008; Rintisch et al., 2014) genotype map of the HXB/BXH panel and convert this map to the latest rat genome assembly (rn6), we genotyped the 30 HXB/BXH recombinant inbred panel lines using a custom designed Affymetrix RATDIV SNP Array at 805,399 variable genetic positions, as described previously (Baud et al., 2014). In short, genotyping was performed according to the Affymetrix SNP chip 6.0 protocol using 250 ng (RNase A-treated) genomic DNA, isolated from rat liver tissue and digested with StyI and NspI, respectively. Genotypes were called and high-quality markers were selected from the 805,399 genotyped SNPs. For this, the original 25-mer Affymetrix probes were first remapped to the latest Ensembl rat genome build (Rnor 6.0) (Flicek et al., 2014) using BLAST (Camacho et al., 2009), requiring the wild type or variant probe to map uniquely within the entire rat genome (as described previously in (Baud et al., 2014)). We furthermore excluded (i) SNPs within 13 base pairs of an indel, (ii) missing or heterozygous variant calls, (iii) monomorphic markers, and (iv) SNPs with a call rate lower than 0.99. The resulting genotype calls could be collapsed into 2,957 genotype blocks, or strain distribution patterns (SDPs), with an average size of 0.75 Mb. Collapsing genotypes into SDPs increased the power for downstream QTL mapping, as not every SNP had to be tested individually. An SDP changed to a next SDP as soon as one of the 30 lines consistently switched genotypes. As some SDPs can occur more than once in the genome, e.g. by chance or because of genotyping or genome assembly errors, we merged such SDPs into a single, globally uniquely occurring SDP, whilst preserving positional information. Subsequently, we merged identical SDPs if separated by a single alternating SDP (e.g. due to a SNP genotyping error). This results in a set of 1,685 unique SDPs that we subsequently used for QTL mapping.

#### Ribosome profiling of heart and liver tissue

For ribosome profiling and mRNA-seq, snap-frozen and powdered tissue was obtained from the animals described in the ‘Animal models’ section. For all samples except for the transgenic *Endog* rats and the *Endog* knockout mice (see below), ribosome profiling was performed using the TruSeq Ribo Profile (Mammalian) Library Prep Kit (Illumina, San Diego, CA; USA), according to a TruSeq Ribo Profile protocol optimized for use on tissue material, as described previously (Heesch et al., 2019; Schafer et al., 2015). In short, ± 50-100 mg powdered tissue was lysed for 10 minutes on ice in 1 mL lysis buffer consisting of 1 × TruSeq Ribo Profile mammalian polysome buffer, 1% Triton X 100, 0.1% NP 40, 1 mM dithiothreitol, 10 U ml^−1^ DNase I, cycloheximide (0.1 mg ml^−1^) and nuclease-free H_2_O. Using immediate repeated pipetting and multiple passes through a syringe with a 21G needle we dissociated tissue clumps to create a homogenous lysate that facilitates quick and equal lysis of the tissue powder. Samples were next centrifuged at 20,000g for 10 minutes at 4°C to pellet cell and tissue debris. Per sample, 400-800*μ*L of lysate was further processed according to the TruSeq Ribo Profile (Mammalian) Reference Guide with the additional modification of 8% PAGE selection directly after PCR amplification of the final library. For all samples, ribosome profiling library size distributions were checked on the Bioanalyzer 2100 using a high sensitivity DNA assay (Agilent; 5067-4626), multiplexed and sequenced on an Illumina HiSeq 2500 producing single end 1×51nt reads. HXB/BXH RI panel samples were always processed in large batches of maximum 30 samples to avoid a sample processing bias.

For heart tissue of transgenic and wild type SHR/Ola rats, as well as *Endog* knockout and wild type C57BL/6 mice, a slightly modified procedure was used due to the termination of the TruSeq RiboProfile kit production by Illumina. The isolation of ribosome footprints is identical to the procedure with the TruSeq kit and as described in (Heesch et al., 2019), except for the use of 7.5uL Ambion RNase 1 (ThermoFisher Scientific AM2295; 100U/uL). Following footprint isolation and PAGE purification, footprints were phosphorylated (NEB T4 PNK; New England Biolabs M0201) and used as input for small RNA library prep using the NEXTflex Small RNA-Seq Kit v3 (Bioo Scientific - PerkinElmer NOVA-5132-06). Libraries were prepared according to manufacturer’s instructions (V19.01), size-selected on 8% PAGE gels (ThermoFisher Scientific EC6215BOX) and QC’d on a Bioanalyzer 2100 (high sensitivity DNA assay; Agilent; 5067-4626). Libraries displayed an average size of 157 bp and were sequenced in a multiplexed manner averaging 4 samples per lane on an Illumina HiSeq 4000. Downstream Ribo-seq data QC shows identical read quality, library complexity and footprint periodicity as libraries generated by Illumina’s TruSeq RiboProfile procedure.

#### Replicate HXB/BXH Ribo-seq experiments

On average, each genomic locus within the HXB/BXH RI panel is shared by 15 animals, as all 30 RI lines are a homozygous mixture of 2 genetic backgrounds (BN-*Lx* and SHR/Ola). To assess the biological variability across individual animals of each HXB/BXH RI line, we performed replicate Ribo-seq experiments on liver tissue of 3 animals (i.e. biological replicates) for 2 of the 30 RI lines: BXH12 and BXH13. For each, we find Pearson correlations > 0.99 across biological replicates, reassuring the high quality of our data and reproducibility of the library preparation and sequencing approach (**Figure S1C**).

#### mRNA-seq and totRNA-seq

For mRNA-seq and totRNA-seq, total RNA was isolated using TRIzol Reagent (Invitrogen; 15596018) using 5-10mg rat and mouse tissue of the exact same powdered tissue samples (from the exact same animals) used for Ribo-seq. RNA was DNase treated and purified using the RNA Clean & Concentrator™-25 kit (Zymo Research; R1018). RIN scores were measured on a BioAnalyzer 2100 using the RNA 6000 Nano assay (Agilent; 5067-1511). Poly(A)-purified mRNA-seq libraries or ribosomal RNA depleted totRNA-seq libraries were generated from the same sample of high quality RNA (average RNA Integrity Number (RIN) for HXB/BXH rats of 9.1 (**Figure S1A**). RNA-seq library preparation was performed according to the TruSeq Stranded mRNA or total RNA Reference Guide, using 500ng of total RNA as input. Libraries were multiplexed and sequenced on an Illumina HiSeq 2500 or 4000 producing paired-end 2×101nt reads.

#### Polysome profiling of congenic rat hearts

Powdered left ventricular heart tissue (3 replicates per congenic line) was lysed in polysome lysis buffer composed of 20 mM Hepes pH 7.5, 5 mM MgCl2, 300 mM KCl, 2 mM DTT, 100 *μ*g/mL cycloheximide, 0.2% NP-40 and 40 U/*μ*L RNAseOut (Invitrogen). Following a 30-minute incubation at 4°C in rotation, the lysed tissue samples were centrifugated for 15 minutes at 20,000 × g at 4°C. An aliquot of the lysate was used to quantify total RNA concentration using the Direct-zol RNA kit (R2051; Zymo, USA) according to the manufacturer’s instructions. From the clear supernatants of the lysates, 15 *μ*g of total RNA was loaded onto 10 – 50% linear sucrose gradients prepared in polysome buffer (20 mM Hepes pH 7.5, 5 mM MgCl2 and 300 mM KCl, 2 mM DTT), and centrifuged at 32,000 rpm (129,311 × g) (SW40Ti rotor, Beckman) for 177 minutes at 4°C. Sucrose gradient fractions were separated using a Biocomp Piston gradient fractionation system associated to a Biorad fraction collector (Biorad model 2110 Fraction Collector) into 42 fractions of 300 *μ*l each, and the absorbance was monitored at 254 nm with an ultraviolet absorbance detector (Biorad model EM-1 Econo UV monitor) to record the polysome profile. Fractions corresponding to the monosomes, light, medium, and heavy polysomes were pooled separately. RNA was extracted with 3x volumes of TriFast-FL (VWR, USA) and purified using Direct-zol RNA kit (Zymo, USA) according to the manufacturer’s instructions. RNA was DNase treated and purified using the RNA Clean & Concentrator™-25 kit (Zymo Research; R1018). RIN scores were measured on a BioAnalyzer 2100 using the RNA 6000 Nano assay (Agilent; 5067-1511). Ribosomal RNA-depleted totRNA-seq libraries were generated from high quality RNA (**Table S1**). RNA-seq library preparation was performed according to the TruSeq Stranded total RNA Reference Guide, using 200ng of total RNA as input. Libraries were multiplexed and sequenced on an Illumina HiSeq 4000 producing paired 2×78nt reads.

#### Sequencing data alignment

Prior to mapping, ribosome-profiling reads were clipped for residual adapter sequences and filtered for mitochondrial, ribosomal RNA and tRNA sequences. Next, we trimmed the 2×101 nt mRNA-seq reads to 29-mers (matching Ribo-seq footprint lengths, which peak at 28-29 nt) and processed those mRNA reads exactly the same as the ribosome profiling data, in order to avoid a downstream mapping or quantification bias due to read length or filtering. For mapping of the HXB/BXH rat RI panel data, we first used Tophat2 v2.1.0 (Kim et al., 2013) to align the full-length 2×101 nt mRNA-seq against the rat reference genome (Rattus Norvegicus rn6, Ensembl release 82), in order to obtain all splicing events naturally occurring in heart and liver tissue. Next, all 29-mer trimmed mRNA and ribosome profiling data were mapped using the splice junction information gathered from the alignment of the full-length mRNA-Seq reads. TopHat2 was used for the initial sequencing data alignment and splice junction determination of the HXB/BXH data analysis, as at the time this project was initiated current state-of-the-art alignment tools were not yet available. Sequencing data was aligned to the reference genome, and not to reconstructed SNP-infused genomes, because the number of allowed mismatches per 29-mer (2 mismatches) suffices to overcome a mapping bias caused by SHR-specific SNPs. We tested this reasoning extensively by aligning replicate trimmed mRNA-seq and Ribo-seq data of SHR/Ola animals (5 replicates) (Schafer et al., 2015) to the BN reference genome or to an SHR/Ola SNP-infused genome. Moreover, we detected no significantly differentially expressed genes i.e. genes for which the expression change could be attributed to a mapping bias driven by local genetic variation. On average, for the HXB/BXH Ribo-seq data we can uniquely align 27.8M Ribo-seq reads for left ventricular tissue samples and 41.5M Ribo-seq reads for liver tissue samples, equaling between 71% and 87% of the total number of sequenced reads used for mapping.

For Ribo-seq and RNA-seq data obtained from congenic rats, transgenic rats, knockout mice and polysome fractionation experiments, sequencing alignment strategies were identical to described above, but using STAR 2-pass v2.7.1a (Dobin et al., 2013) instead of TopHat2 to improve mapping accuracy and speed. Mice data was mapped to the Mus Musculus reference genome mm10, Ensembl release 85. We used STAR to align the previous datasets mapped with Tophat2 and we found Pearson correlations > 0.99 across both methods, supporting the reproducibility of the data regardless of the mapping algorithm. Data QC of all Ribo-seq libraries was performed using Ribo-seQC v1.1 (Calviello et al., 2019).

#### Identifying translated open reading frames

To define the set of translated genes in rat heart and liver, we used RiboTaper v1.3 (Calviello et al., 2016) with standard settings to detect open reading frames that display the characteristic 3-nt codon movement of actively translating ribosomes. For each sample we selected only the read lengths for which at least 70% of the reads matched the primary ORF in a meta-gene analysis. This results in the inclusion of footprints of the most prominent read lengths: 28 and 29 nucleotides. The final list of translation events was stringently filtered requiring the translated gene to have an average mRNA-seq RPKM ≥1 and be detected as translated by RiboTaper in at least 10 out of 30 HXB/BXH RI lines. We did not only retain canonical translation events, but also translated short ORFs (sORFs) detected in long noncoding RNAs (lncRNAs), or upstream ORFs (uORFs) positioned in front of primary ORFs of annotated protein-coding genes. LncRNA sORFs were required to not show sense and in-frame overlap with annotated protein coding genes. We categorically grouped noncoding genes with antisense, lincRNA, and processed transcript biotypes as long noncoding RNAs (lncRNAs), if they matched specific filtering criteria described previously (van Heesch et al., 2019). Upstream ORFs encompass both independently located (non-overlapping) and primary ORF-overlapping translation events. Primary ORF-overlapping uORFs were distinguished from in frame, 5’ extensions of the primary ORF requiring each overlapping uORF to have a translation start site before the start of the canonical CDS, to end within the canonical CDS (prior to the annotated termination codon) and to be translated in a different frame than the primary ORF i.e. to produce a different peptide. We combined both types of uORFs into a single uORF category as we detect no differential impact of each uORF category on the primary ORF TE, in accordance with previous work (Heesch et al., 2019). For the visualization of P-site tracks (**Figure S3C**) we used plots generated by Ribo-seQC (Calviello et al., 2019).

#### Quantifying mRNA expression and translation

Gene-or feature-specific expression quantification was restricted to annotated and identified translated (coding) sequence and performed using HTSeq v0.9.1 (Anders et al., 2015) with default parameters. For quantification of the *Ttn* gene, which codes for the longest protein existing in mammals, we used a custom annotation (Heesch et al., 2019; Schafer et al., 2017) as *Ttn* is not annotated in the current rat gene annotation. For this reason, *Ttn* was initially not included in the QTL mapping analyses, but later on added to define the effect of its length on *Ttn*’s translational efficiency. Moreover, we masked one of the two identical SURF cluster regions in the rat genome (chr3:4,861,753-4,876,317 was masked and chr3:5,459,480-5,459,627 was included), as both regions shared 100% of nucleotide identity and the six expressed SURF genes could not be unambiguously quantified. In parallel to the counting strategy outlined above, for quantifying ribosome association in small and long noncoding RNAs, we additionally ran HTSeq on exonic gene sequences allowing for multiple mapped counts, as many snoRNAs exhibit high sequence homology and they cannot be quantified using only unique counts. In summary, we thus used (i) uniquely mapping CDS-centric counts for mRNA and translational efficiency quantifications, and (ii) multimapping-allowed exonic counts for noncoding RNA quantifications (e.g. *SNORA48*).

The mRNA-seq and Ribo-seq count data was normalized using a joint normalization procedure (DESeq2 v1.26.0 (Love et al., 2014)) as suggested previously (Xiao et al., 2016). This allows for the determination of size-factors for both datasets in a joint manner, as both count matrices follow the same distribution. This is crucial for the comparability of the two sequencing-based measures of gene expression, which for instance becomes important for calculating a gene’s translational efficiency (TE). The TE of a gene can be calculated by taking the ratio of Ribo-seq reads over mRNA-seq reads (Ingolia et al., 2009), or, when biological replicates are available, calculated via specialized DESeq2-based tools (Li et al., 2017; Xiao et al., 2016; Zhong et al., 2017). As we here require sample-specific TE values for downstream genetic association testing with QTL mapping, we regress out the measured mRNA-seq expression from the Ribo-seq expression levels using a linear model. This allows us to derive residuals for each sample-gene pair, that we subsequently subject to QTL mapping. Thus, the TE refers to the residuals of the linear model: resid (lm (normalized_Ribo-seq_read_counts normalized_mRNA-seq_read_counts)). The main advantage of TE values obtained with this calculation is that we retain a quantitative range suitable for QTL mapping, which would not be the case for ratio-based TEs.

#### Pairwise association testing using Matrix eQTL

In order to understand the impact of genetic variants on gene expression regulation, we performed quantitative trait locus (QTL) mapping using the linear regression model-based Matrix eQTL v2.1.1 (Shabalin, 2012). For association testing, non-unique SDPs are grouped and associations of surrounding SDPs are considered when defining the correct SDP location, thereby avoiding falsely assigned distant QTLs because of misplaced contigs in the rn6 rat genome assembly. For this, we reasoned that true associations are likely visible in surrounding SDPs, as genotype changes between two neighboring SDPs are usually gradual, and only a statistically unlikely multitude of recombination events between two neighboring SDPs would fully quench the detected association. After each association is assigned to the correct SDP, we performed a Benjamini-Hochberg correction on local and distant associations separately. A QTL is defined as ‘local’ if it meets an FDR threshold ≤ 0.05 and when it locates within the SDP block of the gene locus for which the association was detected. Similarly, a distant QTL is defined as a trait-associated locus when it meets an FDR threshold ≤ 0.1 and is located on a chromosome different from the one that hosts the associated gene. Subsequently, we performed permutation testing to determine the significance of local and distant associations, by deriving the distribution of test statistics under the null hypothesis that there is no association. We therefore randomized all samples in the gene expression matrix and performed 10,000 runs of Matrix eQTL on the original genotype matrix. A significant association was defined as having an empirical p-value ≤ 0.0015 (less than 15 more extreme p-values in 10,000 permutations). For all types of QTLs tested in this study (eQTLs, riboQTLs, teQTLs and uORF-QTLs), the same association settings and filtering criteria are applied. Throughout the manuscript, QTL numbers reported are gene-centric, i.e. if for instance two neighboring SDPs show significant association with the same gene, a single association is counted. When a given genes associates with both local and distant SDPs, these associations are reported separately. We additionally tested all available technical covariates for a potential impact on our results. These included (i) date of tissue processing, (ii) individual who prepared the libraries, (iii) RIN of the sample, (iv) library concentration (after PCR amplification), (v) date of library PCR and (vi) sequencing batch. None of these technical covariates showed a significant impact on our data (ANOVA p-values between 0.11 and 0.97; using the first PC1 that describes 50-80% of the variance in the data), and were thus not considered during subsequent analyses. All detected significant association results are reported in **Table S2** (eQTLs, riboQTLs and teQTLs) and **Table S3** (uORF-QTLs).

#### Detection of tissue-specific and recurrent QTLs

Gene expression can be regulated in a highly tissue- and cell-type specific manner and genetic effects on mRNA expression can similarly be both specific to, or shared amongst, tissues or cell types (Aguet et al., 2019; GTEx Consortium et al., 2017). Considering only genes expressed in both tissues, both eQTLs and teQTLs show limited recurrence in QTL detection, indicative of high tissue specificity. Even though 83% of genes with cardiac eQTLs (605 out of 727) and 66% of genes with liver eQTLs (248 out of 377) are expressed in both tissues, we could only detect the same eQTL for 126 of these (17%). Similarly, the vast majority genes with teQTLs are expressed in both tissues (88% and 100% in heart and liver, respectively), though only a small fraction of teQTLs (n = 20; 9%) was independently detected in both. All but one of these recurrent eQTLs and teQTLs result from local associations (**Table S2**), indicating strong enrichment of recurrent local over distant QTLs. This is in line with previous observations across human tissues (Aguet et al., 2019; GTEx Consortium et al., 2017) and, in our study, likely influenced by the higher detection sensitivity for local over distant QTLs. A single distant eQTL for *Tmcc2* forms the exception being regulated in *trans* in both tissues (**Table S2**). Although isoform-specific expression regulation of human *TMCC2*, driven by local changes in chromatin dynamics, was previously shown to be of biological importance (Ludwig et al., 2019), its distant control was not yet known.

#### Finding causal variants for local teQTLs

To identify potential causal variants underlying teQTLs, we infused our genotype maps with known SHR/Ola- and BN-*Lx* - specific indels and SNVs that were previously identified through whole-genome sequencing (Atanur et al., 2013; Hermsen et al., 2015; Simonis et al., 2012). Among all genes with a local QTL (either eQTL, riboQTL or teQTL; **Table S2**), we detect only 8 coding sequence variants with a predicted deleterious consequence resulting in one stop gain, one essential splice-site mutation and six missense mutations (Kumar et al., 2009; McLaren et al., 2016). Of these, only a single missense variant in the *Lss* gene is associated with TE in the heart (teQTL p_adj_ = 0.0014; **Table S2**). We find no variants altering the local translation initiation context or Kozak sequence - a previously proposed frequent cause of local teQTLs (Cenik et al., 2015).

#### Detecting distant QTL hotspots with HESS

HESS (Bottolo et al., 2011a) is a generic Bayesian variable selection approach, associated with an evolutionary stochastic search algorithm (Bottolo et al., 2011b), and developed to tackle the challenging integrative task of linking parallel high-dimensional multivariate regressions in a computationally efficient way. When *q* genes are predicted by the same set *p* of SNPs, in HESS the prior probability of association between gene *k* (k = 1,…,*q*) and SNP *j* (j = 1,…,*p*) is decomposed into its marginal effects, i.e., *π*_k,j_ = *π*_k_*xρ*_j_,*π*_k,j_ ∈ [0, 1]. In this formulation, *ρ*_j_ ≥ 0 captures the relative “propensity” for SNP *j* to influence several genes at the same time. The SNP specific “propensity” *ρ*_j_ inflates/deflates the probability *π*_k_ of selecting any SNP to be associated with gene *k* in a multiplicative fashion, i.e., the baseline risk for gene *k* to be associated to any SNP is increased/decreased by the “propensity” of SNP *j* to be a key regulatory marker or “hotspot”. For each gene *k*, the *a priori* baseline risk and the corresponding level of sparsity are controlled through a suitable choice of the hyper-parameters of the density*ρ*(*π*_k_). We ran R2HESS v1.0.1 (Lewin et al., 2016) with default parameters. The marginal posterior probability of inclusion 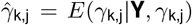 indicates the strength of association between gene *k* and SNP *j* after observing the data **Y** and it is calculated as the number of times a particular gene-SNP pair has been selected. Significant gene-SNP associations were declared using a non-parametric FDR approach, where a mixture model of two beta densities was chosen to model the null H_0_ and the alternative H_1_ distributions. We ran the Expectation-Maximization algorithm (McLachlan & Krishnan, 2008) on 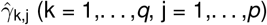 to estimate the parameters of mixture model and, for a fixed FDR level, we calculated the optimal cut-off point on *t* such that the estimated FDR is not greater that the desired one. Finally, the proportion of genes associated with each SNP is defined as the average number of genes that are significantly predicted by each SNP. This measure helps to prioritize SNPs that influence multiple genes at the same time and allows the discovery of so-called regulatory hot-spots, i.e., genetic loci that are associated with a large number of mRNAs.

#### Western blot analysis and quantification

Frozen left ventricle tissues from congenic rats (SHR.BN-(3S) and SHR.BN-(3L)) were lysed in ice-cold modified RIPA buffer (150 mM NaCl, 50 mM Tris HCL pH 7.4, 1% Triton X-100, 0.5% sodium deoxycholate, 0.1% SDS, 5 mM EDTA and 2 mM EDTA) containing protease (cOmplete™, EDTA-free Protease Inhibitor Cocktail) and phosphatase (PhosSTOP) inhibitors as described in (Schiattarella et al., 2019). After incubation on ice for 30 min, samples were centrifuged at 20,000 g for 15 min at 4°C and supernatants were transferred to new pre-chilled tubes. Proteins were denatured for 10 min at 70°C in NuPAGE LDS Sample Buffer (4X) (Invitrogen; NP0007) and NuPAGE Sample Reducing Agent (10X) (Invitrogen; NP0009) and separated on NuPAGE 4-12% Bis-Tris Protein Gels (Invitrogen; NP0343BOX) for 30 min in MES buffer (Invitrogen; NP0002) at 200 V. Gels were blotted on PVDF membranes (Immobilon-PSQ Membrane, Merck Millipore; ISEQ00010) and membranes were stained with the following primary antibodies: p-IRE1*α* Ser724 (NB100-2323, Novus Biological), IRE1*α* (3294, Cell Signaling Technology), XBP1s (83418S, Cell Signaling Technology), GAPDH (ab125247, Abcam), HSP60 (12165, Cell Signaling Technology) and TOM20 (42406, Cell Signaling Technology). Protein expression was measured with chemiluminescence and quantified using Image Studio Lite software (version 5.2, LI-COR). Background-subtracted densitometric signals from SHR.BN-(3S) and SHR.BN-(3L) samples were normalized against the loading control (and the unphosphorylated protein form in case of p-IRE1*α* Ser724) and statistical significant differences between SHR.BN-(3S) and SHR.BN-(3L) samples were determined using unpaired Student’s t-tests.

#### Stoichiometry of the cardiac thin filament

The thin filament is composed of five sarcomere subunits *-Actc1*,*Tpm1*,*Tnnc1*,*Tnnt2*,*Tnni3-* where each unit has a known proportion of 7:1:1:1:1 (Thompson & Metzger, 2014). So as to study how the production rates of the five thin filament proteins deviate from the compositionally stoichiometric optimal ones in the HXB/BXH RI and the congenic rat samples, we estimated the observed proportions by correcting the DESeq2-normalized counts by CDS length and by gene turnover rate. Gene turnover rates for *Actc1*, *Tpm1*, *Tnnc1*, *Tnnt2*, and *Tnni3* have been previously estimated to be 10.3, 5.3, 3.2, 3.5, and 5.5 days, respectively (Martin, 1981).

#### Excluding a technical basis for the length effect

Theoretically, sample-specific gene length biases can artificially induce length-related expression differences that in turn contribute to incorrect enrichment of GO terms related to short (e.g. ribosomal) or long (e.g. ECM) proteins (Mandelboum et al., 2019). However, for multiple reasons we deem it highly unlikely that a technical or analytical bias could be responsible for the length-dependent effect observed in our study. First, the RI lines are all genetic mosaics and the length-dependency is specific for a single locus. Second, the length effect is specific to the heart and absent in liver. Third, data generation, normalization and statistical analysis are all identical for all sequencing samples analyzed. Fourth, no single documented technical covariate explains any of the variance across samples (e.g. date of tissue processing, library preparation batch, sequencing flow cell, or RNA integrity of the sample; see **Table S1**). Fifth, Ribo-seq and polysome fractionation experiments in congenic lines fully reproduce the translation phenotype, indicating a model- and technology-independent effect. Sixth, the effect is absent in RNA-seq data and the correlation with length is stronger for CDS length than for total transcript length. Lastly, previous work on *SNORD24* revealed a highly similar polysome half-mer phenotype accompanied by a length-dependent effect on TE (Thompson & Metzger, 2014).

### QUANTIFICATION & STATISTICAL ANALYSES

The generation of figures and execution of statistical tests were performed using R (R Development Core Team, 2016). GO enrichment analyses were performed using gProfiler2 v0.1.8 (Reimand et al., 2016Reimand et al., 2016). A detailed list of software used for data processing, quantification and analysis is stated in the respective **Methods** sections. We used DESeq2 v1.26.0 (Love et al., 2014) to perform differential gene expression analyses for mRNA-seq and Ribo-seq data. Differentially expressed genes were defined with an FDR ≤ 0.05 and a log2 fold change ≤ 1/1.2 or 1*1.2 for downregulated and upregulated genes respectively. Correlation coefficients between coding sequence (CDS) length and fold changes (FC) in gene expression were based on the Standardized Major Axis (sma) Estimation model (R package ‘smatr’) (Warton et al., 2012). Only CDS with a minimum length of 100 nucleotides and an average number of DESeq2-normalized counts higher than 10 were considered for correlation analyses and plotting. Statistical parameters such as the value of n, mean/median, standard deviation (SD) and significance level are reported in the figures and/or in the figure legends. The “n” represents the number of animals in **Figure 1A**, **Figure S1F** and the **Methods** section “Experimental model and subject details”, or the number of genes with QTLs in **Figure 1E**, the Results section “Identification of translational efficiency QTLs in the HXB/BXH panel” and the **Methods** section “Detection of tissue-specific and recurrent QTLs”. The “p_adj_” represents the FDR significance values calculated by DESeq2 in **Figure 2F**, **Figure 4C** and **Figure 4D**.

## Supplemental Information

### Supplemental Tables and Legends

**Table S1, related to Figure 1 -** Sample information for all sequenced rat and mouse tissue samples, including all open reading frames (ORFs) detected in rat heart and liver. Provided as a separate Excel file.

**Table S2, related to Figure 1 -** Table with local and distant QTL mapping results for rat heart and liver. Includes mRNA expression level QTLs (eQTLs), ribosome occupancy QTLs (riboQTLs) and translational efficiency QTLs (teQTLs). Provided as a separate Excel file.

**Table S3, related to Figure 1 -** Table with upstream ORFs identified in rat heart and liver and detected uORFs-QTLs. Provided as a separate Excel file.

**Table S4, related to Figure 2 -** Table with cardiac QTL hotspots as identified by HESS (see STAR methods). Provided as a separate Excel file.

**Figure S1:**
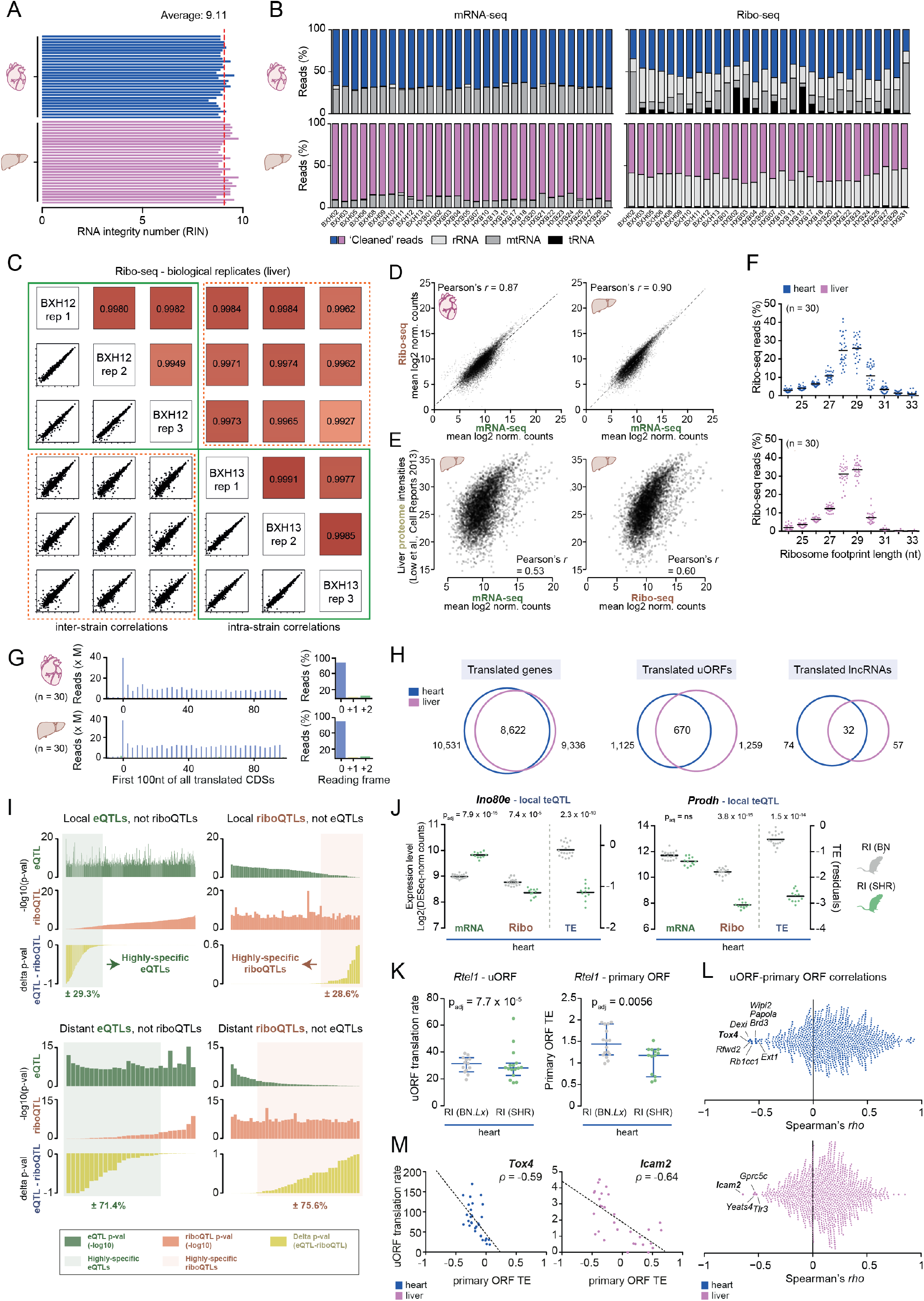
Identification of translational efficiency QTLs in the HXB/BXH panel, related to Figure 1. **(A)** Bar plot with RNA integrity numbers (RIN values) for total RNA isolated from rat heart (top) and liver (bottom). The dashed line indicates the average RIN of 9.11, illustrating the high integrity of the processed tissue samples. **(B)** Stacked bar plots with sequencing read filtering statistics for heart (top) and liver (bottom) tissue. Reads derived from ribosomal RNA (rRNA), mitochondrial RNA (mtRNA) and transfer RNA (tRNA) are removed from the mRNA-seq and Ribo-seq data prior to mapping. This results in a set of ‘cleaned reads’, which are used as input for mapping and downstream data analyses. **(C)** Correlation analyses and scatter plots for biological replicate Ribo-seq data for 3 replicates of two RI lines (BXH12 and BXH13), illustrating the high technical reproducibility of our Ribo-seq approach across biological replicates. **(D)** Correlation scatter plots for heart (left) and liver (right) tissue, showing the correlation between mean mRNA-seq and Ribo-seq based quantifications of gene expression. **(E)** Correlation scatter plots between mRNA-seq or Ribo-seq reads and rat liver proteomics data, as quantified using iBAQ values obtained from MaxQuant (Cox & Mann, 2008). Ribo-seq is a slightly better proxy for final protein levels than mRNA-seq (Pearson’s *r* = 0.60 vs 0.53). **(F)** Dot plot showing the ribosome footprint (Ribo-seq read) length distribution across the 30 lines in rat heart (left) and liver (right). **(G)** Venn diagrams with tissue-specific comparisons of all identified translated genes, translated lncRNAs and translated uORFs. **(H)** Bar plot with a meta-analysis of stacked P-sites derived from Ribo-seq reads in heart (top) and liver (bottom) tissue. Blue bars indicate the number (left) and percentage (right) of footprints that precisely match annotated protein-coding gene open reading frames (ORFs; ± 90%). **(I)** Bar plots with significance values for detected local (top) and distant (bottom) eQTLs and riboQTLs, sorted by the delta of the p-values for both quantitative traits (bottom track of each panel). This analysis illustrates that most eQTLs are prolonged during translation, though they may sometimes near-miss the significance cutoff. Concordant with the teQTL results, this analysis furthermore highlights a highly translation-specific set of QTLs. **(J)** Dot plots with expression values for 2 genes with a highly specific local teQTL in rat hearts. Bars indicate mean values. **(K)** *Rtel1* is the only gene for which a local teQTL and uORF-QTL coincide, though both associations occur with similar directionality (lower translation associate with the SHR/Ola genotype). **(L)** Bee swarm dot plots with correlation values (Spearman’s *rho*) between the translation rates of uORFs and the primary ORF TE in heart and liver. Genes with uORFs that show a strong negative correlation with primary ORF translation are highlighted in red. Overall, most uORFs seem to positively correlate with primary ORF TE, as previously reported for the human heart (Heesch et al., 2019). **(M)** Two rare examples of genes with strongly anti-correlating translation rates for the uORF vs. the primary ORF.

**Figure S2:**
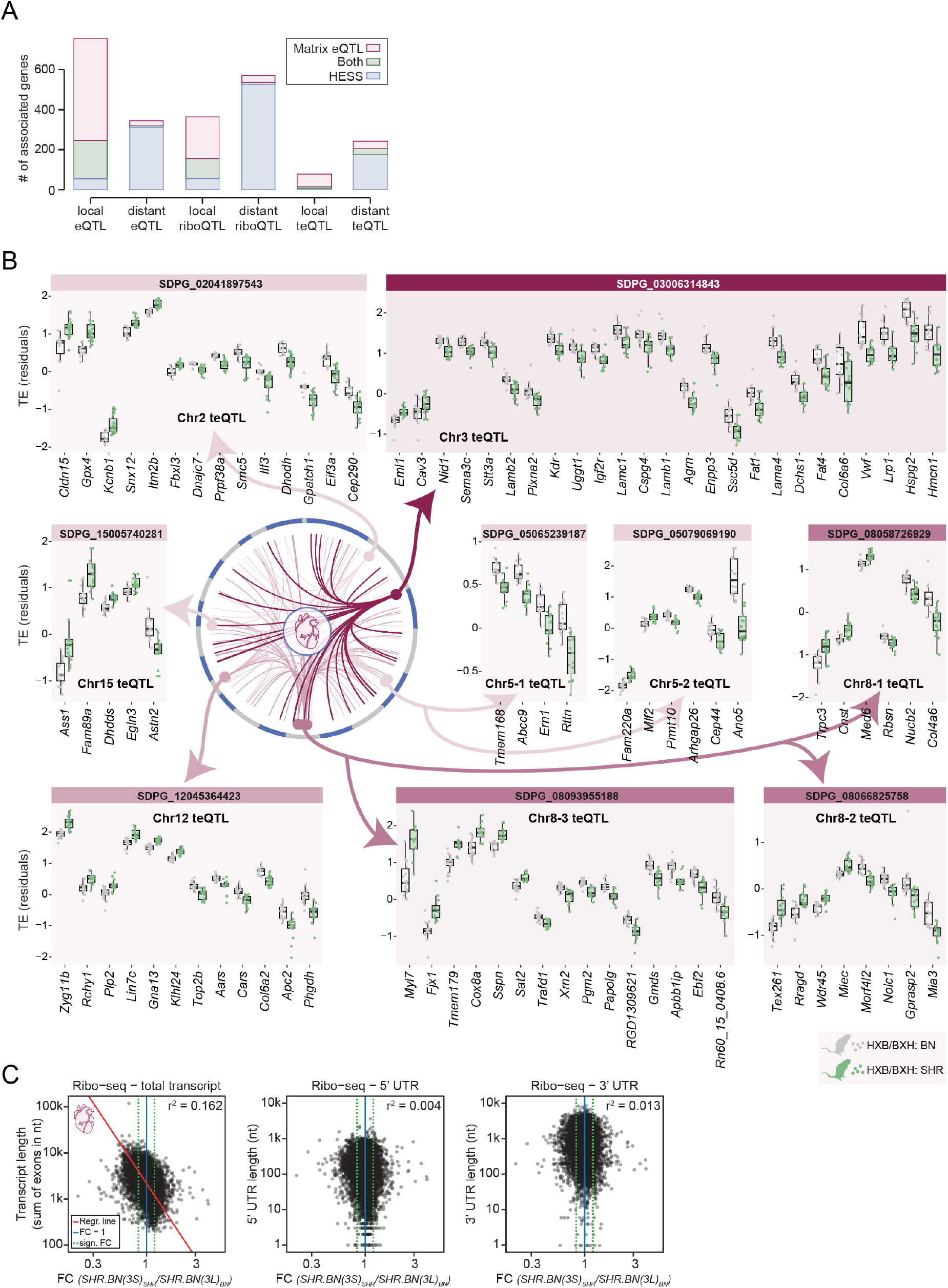
The chromosome 3p teQTL regulates cardiac translation in a protein length-dependent manner, related to Figure 2. **(A)** Stacked bar plot showing the number of associated genes (i.e. with specific QTLs) in the rat heart, for Matrix eQTL, HESS or both. HESS is especially powerful for the detection of distant associations where a single locus can be linked to multiple genes. **(B)** Circos plot with all distant master regulatory teQTLs that associate with the TE of at least 5 genes (as in **Figure 2A**). For each identified distant QTL hotspot, gene-specific TE dot box plots are given for all associated genes. The teQTL coordinates, i.e. the genotype block linked to each set of genes, can be derived from the SDP ID (e.g. SDPG_03006314843 starts at Rat rn6 Chr3:6,314,843). Per teQTL, genes are ordered by effect size and directionality. See also **Table S4**. **(C)** Scatter plots and square correlation coefficients (r^2^) based on standardized major axis (SMA) values between total transcript length, 5’ UTR length, or 3’ UTR length versus the fold change (FC) in gene expression, as measured by Ribo-seq in congenic rat hearts. Obtained correlation coefficients (r^2^) are lower than that of the comparison between CDS length and FC in translation (see **Figure 2E**), indicating that CDS length is the main determinant of the translational efficiency phenotype. For UTR length versus FC in gene expression, only cases with at least 10 DESeq2-normalized counts in both SHR.BN-(3L) and SHR.BN-(3S) rats are displayed. For total transcript length versus FC in gene expression, the correlation is significant (p-value < 2.2 × 10^−16^; Test of correlation coefficient against zero) and the linear model based on fitted SMA method is displayed as a red line.

**Figure S3:**
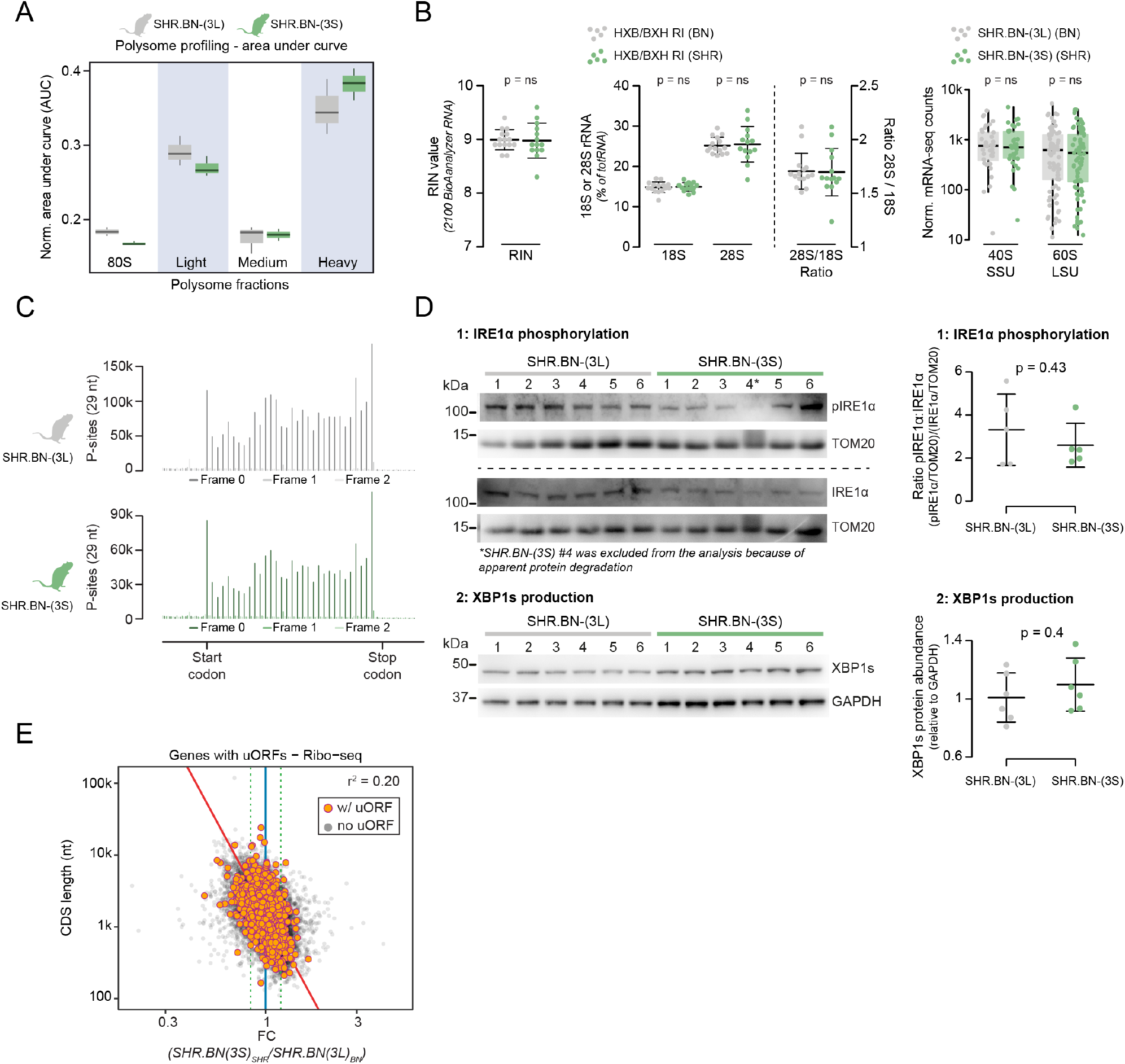
Impaired ribosome assembly and half-mer formation drive the chromosome 3p teQTL, related to Figure 3. **(A)** Rat congenic line comparison of differences in polysomal configuration as measured by normalized area under curves (AUCs) of the polysome profiles, for SHR.BN-(3L) (grey) and SHR.BN-(3S) (green). **(B)** Left panel: dot plots with RIN values of HXB/BXH RI lines separated by Chr. 3p teQTL genotype. RIN values shown are calculated on the Agilent BioAnalyzer 2100 using the RNA nano assay. The middle panel contains dot plots of the HXB/BXH RI lines, with (i) 18S abundance, (ii) 28S abundance and (iii) 28S / 18S ratios, as calculated by the percentage of total RNA with an Agilent BioAnalyzer 2100 RNA nano assay. The right panel contains dot box plots comparing congenic rat line mean mRNA expression values for ribosomal protein genes involved in the structure of the 40S (SSU) and 60S (LSU) ribosomal subunits. Values are separated by local genotype at the chromosome 3p teQTL locus and only left ventricular heart tissue RNA data is shown. Error bars indicate mean values with standard deviation (SD). All these analyses indicate no imbalance between the production levels of both ribosomal subunits. **(C)** Meta-gene codon 3-nt periodicity bar plots displaying P-sites of 29-nt ribosome footprints to illustrate the similarity in translation elongation rates between both congenic lines. The visualized data are a merger of all replicate SHR.BN-(3L) (grey, top) and SHR.BN-(3S) (green, bottom) Ribo-seq 29-nt footprints. Plots are generated with Ribo-seQC (Calviello et al., 2019) and modified to only display the following sections of genes: (i) 25nt before and 33nt after the start codon, followed by (ii) 33nt from the middle of the CDS, and finally (iii) 33nt before and 25nt after the stop codon (Calviello et al., 2019). **(D)** Western blots and dot plots with quantification of band intensities for the ER stress markers IRE1-alha phosphorylation (normalized for IRE1-alpha and TOM20 expression; top) and XBP1s production (normalized for GAPDH expression; bottom). Error bars indicate mean values with SD. These protein level analyses show no difference in the expression or activation of typical ER stress response markers. **(E)** Scatter plot showing CDS length versus fold change (FC (SHR.BN-(3S) vs SHR.BN-(3L)) for Ribo-seq data, highlighting all genes with actively translated uORFs in the rat heart. The square correlation coefficient (r^2^) based on standardized major axis (SMA) is calculated using expression values of this subset of uORF-containing genes only, and precisely matches that of the whole translatome (r^2^ = 0.20), showing that there is no differential regulation of genes with uORFs as opposed to the full set of translated genes.

**Figure S4:**
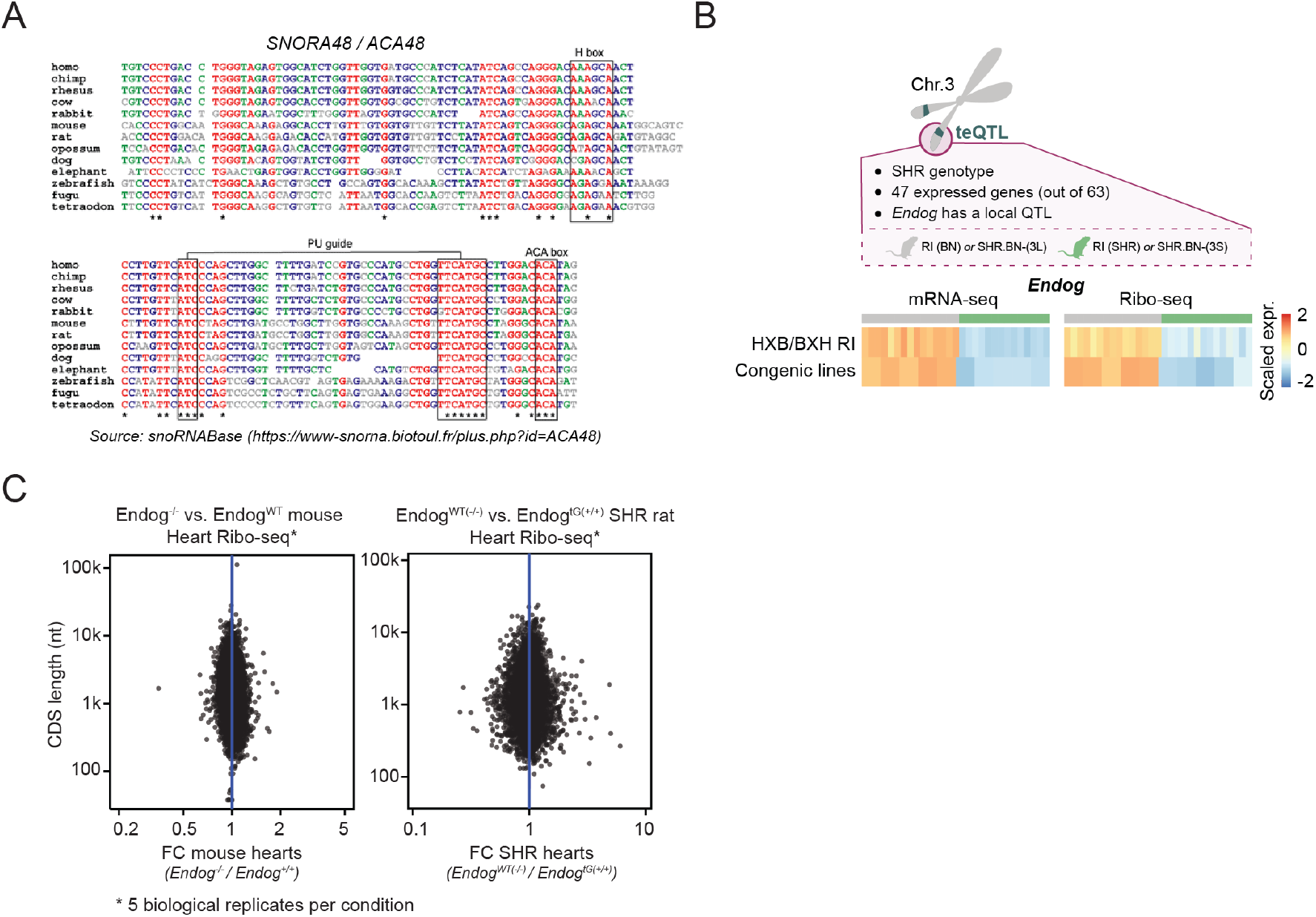
Reduced de novo translation rates reinforce a pre-existing length-bias in TE. **(A)** Multi-species alignment for the H/ACA box snoRNA *SNORA48*, highlighting functional domains (H box, pseudouridylation guide, ACA box), as obtained from snoRNABase v3 (snoRNA-LBME-db; visited: March 2020) (Lestrade & Weber, 2006). **(B)** Schematic visualization of the rat chromosome 3 teQTL and a summary of the expressed genes within this region. Expression values for *Endog* are given. Heatmaps show scaled and normalized expression values. **(C)** Scatter plots for CDS length versus the fold change (FC) in gene expression as measured by Ribo-seq in *Endog*^−/−^ mice vs WT mice (left) and transgenic SHR/Ola rats with partially rescued *Endog* expression versus wild type SHR/Ola (right). Ribo-seq was performed on 5 hearts per condition (see also **Table S1**), though reveals no correlation between CDS length and translation suggesting that *Endog* knockout or transgenic rescue alone is not sufficient to induce or ameliorate the CDS length-dependent shift in TE.

